# Open-source, Hardware-Independent GPU Acceleration for Scalable Nanopore Basecalling with Slorado and Openfish

**DOI:** 10.64898/2026.03.25.714356

**Authors:** Bonson Wong, Gagandeep Singh, Haris Javaid, Kristof Denolf, Kisaru Liyanage, Hiruna Samarakoon, Ira W. Deveson, Hasindu Gamaarachchi

## Abstract

Nanopore sequencing technologies are used widely in genomics research and their adoption continues to accelerate. ‘Basecalling’ is an essential step in the nanopore sequencing workflow, during which raw electrical signals are translated into nucleotide sequences. The current state-of-the-art basecaller, Oxford Nanopore Technologies (ONT) software ‘Dorado,’ relies on proprietary, platform-specific NVIDIA GPU optimisations bundled in the closed-source ‘Koi’ library. As a result, practical, high-speed basecalling is effectively restricted to a narrow class of supported hardware, limiting accessibility, portability, and innovation. We present (1) ‘Openfish,’ an open-source GPU-accelerated nanopore basecaller decoding library that provides a competitive alternative to ONT’s proprietary Koi library; and (2) Slorado, a fully open-source basecalling framework that supports both DNA and RNA with equivalent accuracy to Dorado. Together, Openfish and Slorado remove the hardware lock-in that currently limits high-performance nanopore basecalling. Our framework scales efficiently across heterogeneous computing environments, from low-power embedded devices to GPU-equipped datacenters, without sacrificing speed or accuracy. Openfish and Slorado are available as free open-source packages for basecalling research, optimisation and deployment beyond the constraints of proprietary software and hardware ecosystems: Openfish: https://github.com/warp9seq/openfish, Slorado: https://github.com/BonsonW/slorado.

## 1. Introduction

Nanopore sequencing technologies has become a widely adopted platform in genomics due to their portability, flexibility, cost-effectiveness and ability to generate long DNA or RNA reads [1–4]. Unlike other sequencing technologies, nanopore devices produce continuous electrical time-series ‘raw signal’ data that must be translated into nucleotide sequences through the process of basecalling [5–7]. As the first computational step in a nanopore workflow, basecalling is both computationally demanding and algorithmically complex [6–17]. Recent advances in deep learning have substantially improved basecalling accuracy, with modern basecallers relying on neural network architectures trained directly on raw signal data [6, 15–26].

Oxford Nanopore Technologies (ONT), the primary manufacturer of nanopore sequencing devices, currently provides the state-of-the-art production basecaller, *Dorado* [27]. ONT basecalling models are divided into three categories: fast (FAST), high-accuracy (HAC), and super-accuracy (SUP). FAST models provide the fastest inference with the lowest accuracy; HAC models balance speed and accuracy; and SUP models maximise accuracy at substantially higher computational cost [28]. Version 5 basecalling models from ONT (v5) employ long short-term memory (LSTM)-based recurrent neural networks for FAST and HAC, and attention-based Transformer architectures for SUP.

Dorado’s basecalling pipeline consists of two major steps (Fig. 1A). The first step, *inference*, scores at each time step the transition probabilities between all possible k-mers (subsequences of length k) based on the raw electrical signal [9]. The second step, *decoding*, interprets these probabilities to recover the most likely nucleotide sequence, typically by searching for the optimal path through the time-series of predicted states [9, 13, 21]. To achieve practical runtimes, key components of the inference and decoding steps have been heavily optimised for execution on specific NVIDIA GPUs.

**Fig. 1:**
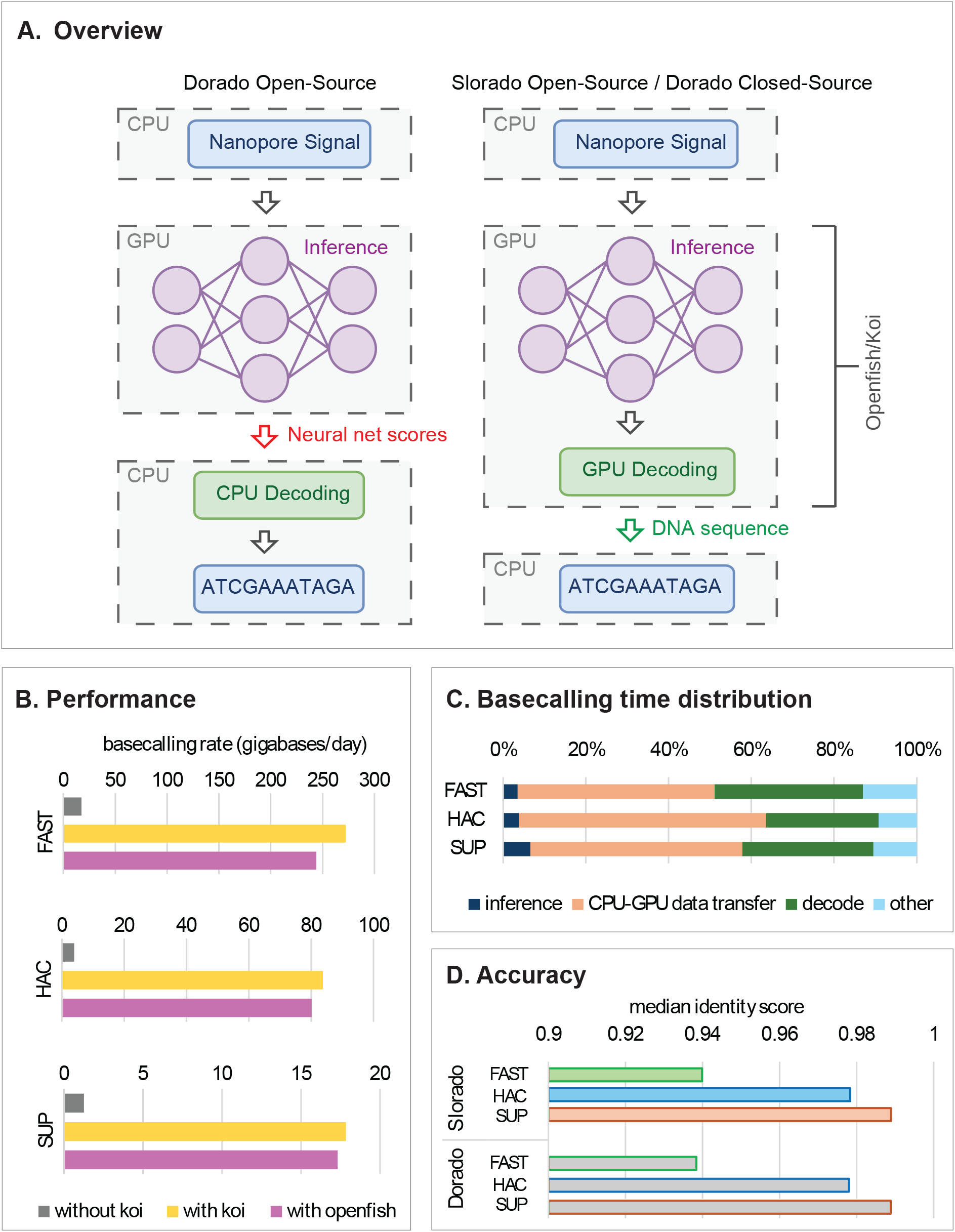
**A**. Overview of Dorado and Slorado basecalling pipelines. Dorado’s open-source execution path performs decoding on the CPU (left panel), which requires the transfer of neural network scores between GPU and CPU (extremely slow). Slorado avoids this by using our open-source Openfish, which implements decoding directly on the GPU, whereas Dorado relies on the closed-source Koi for the same purpose (right panel). **B**. Basecalling rate (the higher the better) when decoding is performed on the CPU without Koi (grey bars), or on the GPU using either Koi (yellow bars) or Openfish (pink bars). **C**. Basecalling time breakdown when using the CPU decoder. **D**. Accuracy of Slorado v0.4.0-beta (with Openfish decoding) against Dorado v1.1.1 (with Koi decoding) on 4*×*NVIDIA A100 GPUs for v5.0.0 DNA basecalling models.

Although the Dorado code is open-source, these critical platform-specific optimisations are bundled in a closed-source, precompiled library called *Koi*. The Koi library implements highly accelerated NVIDIA GPU kernels for the entire decoding stage of basecalling and selected neural layers within the inference step (Fig. 1A). In the absence of Koi, basecalling performance degrades to impractical levels, as we demonstrate in Results. Thus, state-of-the-art ONT basecalling is effectively inaccessible to users working outside the narrow range of computing environments to which ONT has tailored their software. This dependence on proprietary platform-specific acceleration creates a substantial barrier to portability, reproducibility, and broader deployment across heterogeneous computing environments.

Although Dorado includes support for Apple Silicon, the basecalling performance on these systems remains impractical for real-world use, leaving users to rely solely on the newest high-performance and often costly NVIDIA GPUs [29]. It also prevents the adaptation of basecalling software to miniature, low-cost, low-power hardware platforms that are essential to fully realise the potential of nanopore technology for decentralised and portable genomics applications [9, 13, 30, 31].

To address these limitations, we developed *Openfish*, an open-source GPU-accelerated decoding library for modern state-of-the-art basecalling models. Openfish provides performance competitive with ONT’s closed-source Koi, across both LSTM-based recurrent neural networks (used in ONT’s v5 FAST and HAC models) and attention-based Transformer neural networks (used in v5 SUP models). We integrated Openfish into a streamlined, fully open-source basecalling framework called *Slorado*, designed for both end-users and researchers. Slorado enables basecalling at practical speeds on a wide range of heterogeneous GPUs, including both NVIDIA and AMD hardware, while maintaining accuracy equivalent to Dorado. Together, Openfish and Slorado remove proprietary hardware constraints from high-performance nanopore basecalling, expanding accessibility, enabling portable deployment, and supporting transparent, community-driven development and research of next-generation basecalling algorithms.

## 2. Results

### 2.1. Dependence of Dorado on Koi

We first evaluated the impact of removing Dorado’s closed-source Koi component on basecalling performance. Such a configuration represents the fully open-source execution path in Dorado (Fig. 1A; see Methods), where model inference is performed on the GPU via the LibTorch API (the C++ backend of PyTorch), and decoding is performed on the CPU. Without Koi, basecalling a small 0.1 gigabase dataset using an NVIDIA A100 GPU and a 32-core CPU took ∼7 minutes ( ∼17 gigabases/day) for FAST, ∼30 minutes ( ∼4 gigabases/day) for HAC and ∼2 hours ( ∼1 gigabases/day) for SUP (grey bars in Fig. 1B, Supplementary Table S1). In comparison, when Koi was used for decoding, the same dataset was basecalled in only ∼30*×* seconds ( ∼272 gigabases/day) for FAST, ∼2 minutes for HAC ( ∼84 gigabases/day) and ∼7 minutes ( ∼18 gigabases/day) for SUP (yellow bars in Fig. 1B, Supplementary Table S1). This represents a >14*×* slowdown if Koi is not used for decoding (Fig. 1B). When extrapolated to a typical ONT PromethION dataset with ∼100 gigabases (a human genome at ∼30× depth), basecalling without Koi would take ∼5 days for FAST, ∼3 weeks for HAC and ∼3 months for SUP. Thus, without the closed-source Koi package, the open-source Dorado implementation cannot be used to basecall a typical dataset within a practical timeframe.

To investigate the reason for such a large decrease in performance when not using Koi, we benchmarked the time for each major step of the basecalling pipeline (Fig. 1C, Supplementary Table S2). This experiment revealed that the biggest bottleneck lies in the process of copying data from the GPU to the CPU, which consumed ∼50% of the total basecalling time, followed by the actual decode step, which consumed ∼30% of the time. Due to the large amount of data generated from the inference step (e.g., ∼100 GiB for FAST, ∼400 GiB for HAC and ∼1.6 TiB for SUP, for a small 0.1 gigabase dataset), the throughput of data copying between the GPU (where the score data resides) and the CPU (where the decoding algorithm is implemented) becomes the bottleneck.

### 2.2. Openfish and Slorado

To address the bottleneck identified above, we accelerated the decoding algorithm on the GPU, thereby completely bypassing the large amount of copying required from the GPU to the CPU (Fig. 1A; see Methods). We call our implementation Openfish. We benchmarked our Openfish decoder using the same small 0.1 gigabase dataset above. Basecalling performance was similar when using Openfish or the Koi decoder, with Koi being faster by just ∼6% on average (pink bars in Fig. 1B, Supplementary Table S1). Openfish, therefore, has the potential to enable ONT basecalling at practical speeds without the closed-source optimisations in Koi.

To achieve this, we developed Slorado, a fully open-source software for ONT basecalling. Slorado performs basecalling using the same deep learning-based models as Dorado but differs in its software architecture (see Methods). Two key differences are that Slorado takes input signal data in binary SLOW5 (BLOW5) format [32], rather than ONT’s native POD5 [33], and uses Openfish optimisations rather than Koi. Slorado invokes the Openfish library for decoding directly after inference within the GPU itself. The Openfish library takes the direct output of the inference step and returns the resulting nucleobase sequence, quality scores and move table.

The GPU-accelerated portion of Openfish (conditional random field-connectionist temporal classification [CRF-CTC] decoding [9, 34, 35]) is implemented in both programming languages CUDA C and HIP C, supporting NVIDIA and AMD platforms, respectively. In addition, we have integrated several fused layers in Slorado’s attention-based inference step. This includes the fused root mean squared normalisation layers supported by LibTorch, as well as a custom fused implementation of the rotary embedding layer we include as part of the Openfish library. All fused layers are supported in both NVIDIA and AMD, and packaged within Slorado.

### 2.3. Slorado basecalling accuracy

Because Slorado uses the same basecalling models as Dorado, it is expected to deliver basecalled reads with equivalent accuracy. To confirm this, we basecalled a complete HG002 PromethION dataset (HG002-dna in Table 1) with Slorado and Dorado. We measured the accuracy of each basecaller by aligning the reads to the hg38 reference and taking the median identity scores (see Methods).

**Table 1.** The details of the datasets used for the experiments. (estimated gigabases) = (total signal samples) / (sample rate) * (translocation rate).

| Dataset | Description | N reads (millions) | Total signal samples (billions) | Estimated gigabases |
| --- | --- | --- | --- | --- |
| HG002-dna | HG002; PromethION LSK114; 5kHz sample rate; 400 bases/s | 16.1 | 903.2 | 72.25893893 |
| HG002-dna-500k | 500,000 read subset of HG002-dna | 0.5 | 28.1 | 2.24860235 |
| HG002-dna-20k | 20,000 read subset of HG002-dna | 0.02 | 1.1 | 0.08968156 |
| HG002-dna-40× | HG002; PromethION LSK114; 5kHz sample rate; 400 bases/s | 7.2 | 1619.7 | 129.5790768 |
| UHRR-rna | Universal human RNA; PromethION RNA004; 4kHz sample rate; 130 bases/s | 16.4 | 641.1 | 20.83625010 |

We observed very similar accuracy between Slorado and Dorado for all the basecalling models (Fig. 1D, Supplementary Table S3). Slorado exhibited median identity scores slightly higher than Dorado, for example, 0.988923 vs. 0.988889 for SUP. Such minor variations are expected due to differences in floating point precision and GPU library version differences, and are even observed when running the same version of Dorado on different hardware (Supplementary Table S3 and also shown previously in [36]). These results confirm that basecalling outcomes are effectively equivalent between Slorado and Dorado.

We then assessed how Slorado and Dorado affect downstream small variant calling. To do this, we used Clair3 [37] to call variants and evaluated the accuracy using RTG Tools (see Methods). Both Slorado and Dorado achieved a precision and sensitivity of 0.998 for single nucleotide variant (SNV) calling. For short indels (non-SNV), Slorado achieved a slightly higher precision (0.951) than Dorado (0.950), but a slightly lower sensitivity (0.865 vs. 0.870). We then restricted the variant evaluation to homopolymer regions (see Methods), which are known to be challenging for ONT sequencing. Both Slorado and Dorado achieved identical SNV-calling performance, with a precision of 0.986 and a sensitivity of 0.984. For short indels, Slorado achieved slightly higher precision than Dorado (0.860 vs. 0.856), but slightly lower sensitivity (0.720 vs. 0.732). The ROC curves for Slorado and Dorado displayed a near identical overlap across all evaluations (Supplementary Fig. S1).

### 2.4. Comparison with Dorado on matched hardware

We next compared the speed of basecalling with Slorado to Dorado, on matched hardware. Because Dorado incorporates extensive platform-specific neural network optimisations through the Koi library, Slorado was not expected to exceed Dorado’s performance. Our goal was to achieve broadly comparable runtimes, to demonstrate the feasibility of using Slorado as a fully open-source alternative for ONT basecalling.

Since both basecallers support NVIDIA data centre GPUs, we used Australia’s National Computational Infrastructure (NCI) ‘Gadi’ supercomputer, which hosts nodes with 8*×*NVIDIA A100 GPUs, and nodes with 4*×*H200 GPUs. Basecalling was run on the same HG002 PromethION dataset used in previous experiments (HG002-dna in Table 1). On an 8*×*A100 GPU node (hpc-nci-1 in Table 2), Slorado completed basecalling in 1.5 hours for FAST, 2.7 hours for HAC, and 7.6 hours for SUP (Fig. 2A). On a 4*×*H200 GPU node (hpc-nci-2 in Table 2), Slorado completed basecalling in 0.8, 1.6, and 7.4 hours using FAST, HAC, and SUP models, respectively. Dorado was 2-3*×* faster than Slorado when using the SUP model, reflecting the high level of optimisation for this Transformer model. However, Slorado achieved similar speeds when using FAST or HAC models, and even outperformed Dorado when using FAST on the 8*×*A100 GPU node and HAC on the 4*×*H200 GPU node. This surprising observation is explained by the choice of file format. Despite Dorado’s neural network optimisations, on these systems, with a large enough dataset, the basecaller is bottlenecked by the POD5 file format, which has relatively poor file-reading performance in high-performance cluster file systems (e.g., Lustre), as we previously reported [38]. On a custom server with a file system in which POD5 I/O no longer bottlenecks Dorado (server-1 in Table 2), we demonstrate this by running the same experiment (Fig. 2A). Overall, these results demonstrate Slorado’s practical basecalling speeds on systems with NVIDIA data centre GPUs, while providing a fully open-source alternative to Dorado.

**Table 2.** The details of the computer systems used for the experiments.

| System name | n × GPU | MEM /GPU (GB) | n × CPU | total cores, threads | RAM (GB) | Disk/ file system | O/S | Lib Torch | GPU toolkit |
| --- | --- | --- | --- | --- | --- | --- | --- | --- | --- |
| server-1 | 4 × NVIDIA A100 | 40 | 2 × AMD EPYC 7313 | 32, 64 | 540 | 12×HDD (RAID10), ext4(NFS) | Ubuntu 22.04 | 2.9 | CUDA 11.8 |
| hpc-nci-1 | 8 × NVIDIA A100 | 80 | 2 × AMD EPYC 7742 | 128, 256 | 2048 | Lustre | Rocky Linux 8.10 | 2.9 | CUDA 12.8 |
| hpc-nci-2 | 4 × NVIDIA H200 | 141 | 2 × Intel Xeon Gold 6542Y | 48, 96 | 1024 | Lustre | Rocky Linux 8.10 | 2.9 | CUDA 12.8 |
| hpc-pawsey-1 | 4 (8 GCD) × AMD MI250X | 128 | 1 × AMD EPYC 7A53 | 64, 128 | 230 | Lustre | SUSE Linux 15 SP6 | 2.9 | ROCm 6.4 |
| hpc-amd-1 | 8 × AMD MI300X | 192 | 2 × AMD EPYC 9684X | 192, 384 | 2300 | WekaFS | Rocky Linux 9.6 | 2.9 | ROCm 6.4 |
| desktop-1 | 1 × AMD RX 7900XTX | 24 | 1 × Ryzen 7 7700X | 8, 16 | 64 | SSD, ext4 | Ubuntu 22.04 | 2.9 | ROCm 6.3 |
| edge-1 | 1 × Jetson AGX Xavier | 16 | 1 × ARMv8.2 Carmel | 8, 8 | 16 | SSD, ext4 | Ubuntu 20.04 | 2.0 | JetPack 5.1.0, CUDA 11.4 |
| edge-2 | 1 × AMD 760M | 32 | 1 × AMD Ryzen 5 7640HS | 6, 12 | 32 | SSD, ext4 | Ubuntu 22.04 | 2.9 | ROCm 6.2.4 |

**Fig. 2:**
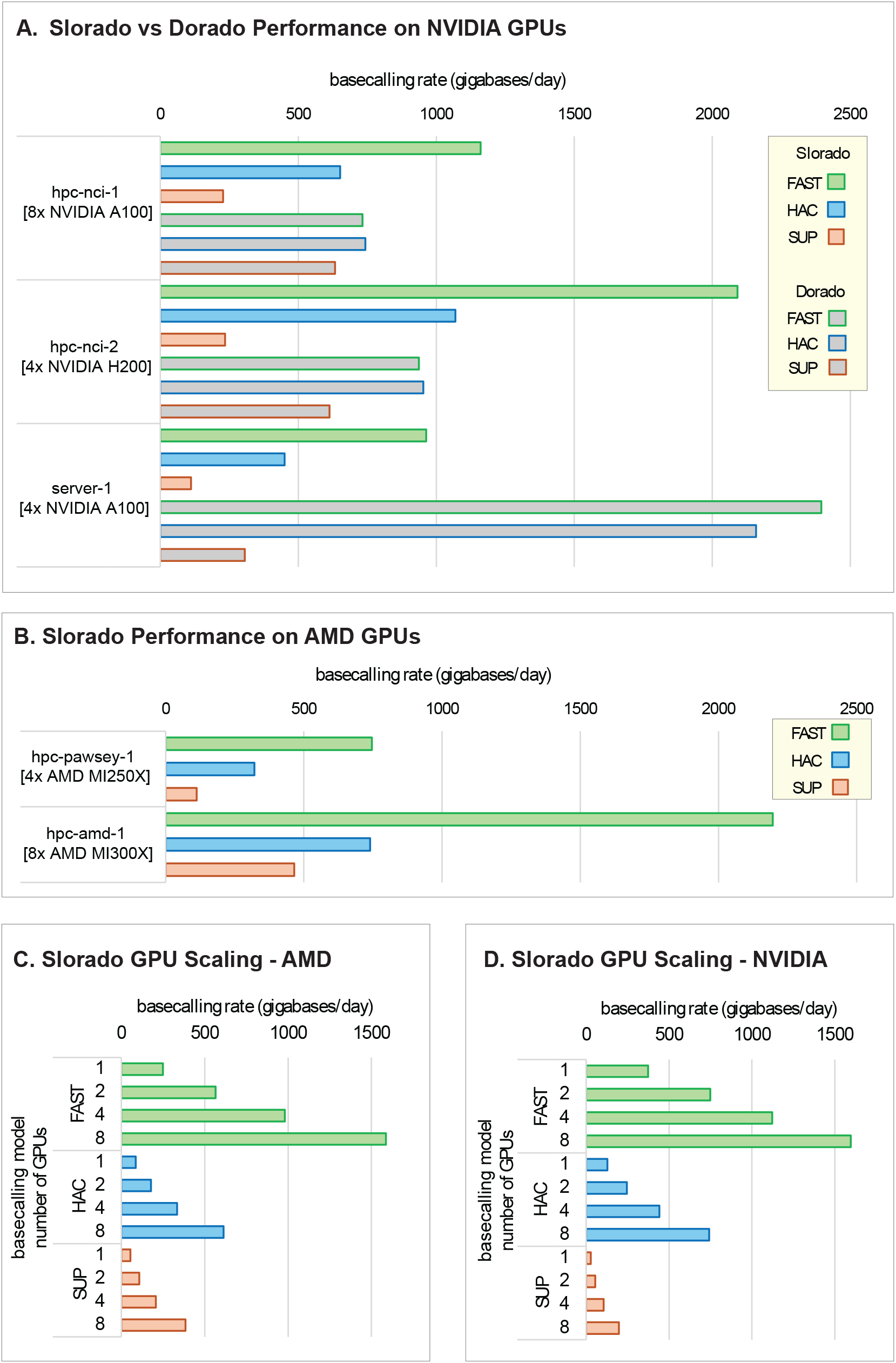
**A**. DNA basecalling rate of Dorado vs. Slorado (the higher the better) on systems with NVIDIA data centre GPUs when using ONT’s v5.0.0 models. Calculated from the total time taken to basecall the HG002-dna dataset. **B**. DNA basecalling rate of Slorado on systems with AMD Instinct™ GPUs when using ONT’s v5.0.0 models. Calculated from the total time taken to basecall the HG002-dna dataset. **C**. Performance scaling of Slorado on a multi-GPU node, when using 1-8 AMD MI300X GPUs. **D**. Performance scaling of Slorado on a multi-GPU node, when using 1-8 NVIDIA A100 GPUs.

### 2.5. Compatibility with AMD GPUs

Our primary goal in developing Slorado was to enable basecalling on GPUs not currently supported by Dorado. Therefore, we next evaluated Slorado basecalling on GPUs manufactured by AMD, rather than NVIDIA. We basecalled the same dataset as above, this time on Australia’s ‘Pawsey’ supercomputer, which is equipped with data-centre class AMD Instinct™ MI250X GPUs. On a single node with 4*×*AMD MI250X GPUs (total 8 Graphics Compute Dies [GCD], hpc-pawsey-1 in Table 2), Slorado completed the basecalling for the complete HG002 PromethION dataset (HG002-dna in Table 1) in 2.3 hours (FAST), 5.4 hours (HAC), and 15.4 hours (SUP), corresponding to throughputs of 746, 321, and 113 gigabases/day, respectively (Fig. 2B, Supplementary Table S3).

On a node with 8*×*AMD MI300X GPUs in the AMD AI & HPC Cluster (hpc-amd-1 in Table 2), the same experiment took 0.8 hours (FAST), 2.3 hours (HAC), and 3.7 hours (SUP), corresponding to 2196, 740, and 465 gigabases/day, respectively (Fig. 2B, Supplementary Table S3). This demonstrates the capacity of Slorado to execute basecalling within practical timeframes on AMD GPUs, which is not supported at all by Dorado.

In doing so, Slorado enables ONT users to harness additional untapped computational resources. For example, the HPC ‘Pawsey’ (one of Australia’s largest academic supercomputers) lacks NVIDIA GPUs and is instead equipped with 192 GPU nodes, each with 4*×*AMD MI250X GPUs, which are currently inaccessible for ONT basecalling.

To evaluate scalability across multiple GPUs, we first measured the speedup in basecalling rate when using 1, 2, 4 and 8 MI300X GPUs on a single node (Fig. 2C). With the FAST model, we were able to scale from 250 gigabases/day on 1 GPU to 1587 gigabases/day on 8 GPUs (6.4*×* relative speedup). With HAC, Slorado’s basecalling rate increased from 86 to 613 gigabases/day (7.2*×* relative speedup), and with SUP, the basecalling rate increased from 54 to 385 gigabases/day (7.1*×* relative speedup). Comparable scaling results were obtained separately for NVIDIA GPUs (Fig. 2D). Therefore, although some time is lost to load imbalance, due to the synchronous pipeline of Slorado’s basecalling framework, Slorado’s basecalling speed roughly scaled with the number of GPUs available.

Next, we evaluated large-scale parallelisation across multiple compute nodes to maximise utilisation of Pawsey’s GPU resources. By partitioning the basecalling tasks into 33 separate parallelisable jobs, Slorado was able to complete SUP basecalling on the complete HG002 PromethION dataset in just 1.2 hours on the Pawsey HPC (Fig. 3A). In this configuration, the first 19 basecalling jobs were executed in parallel (Fig. 3A) before the next 14 jobs were executed in parallel, where each job allocates its own separate node to be run. This was much faster than basecalling on a single node on Pawsey (15 hours for the same dataset; 12.5-fold improvement) and illustrates the potential value of untapped compute resources for ONT basecalling. While the speedup observed here is achieved by allocating a larger number of nodes in parallel, this strategy is well suited for time-critical applications such as ultra-rapid critical care sequencing, where large-scale cloud GPU deployments have previously enabled diagnostic results to be generated within hours [39]. Slorado extends this capability to AMD GPU resources, enabling comparable large-scale deployments and making such computational capacity accessible in environments where it was previously unavailable.

**Fig. 3:**
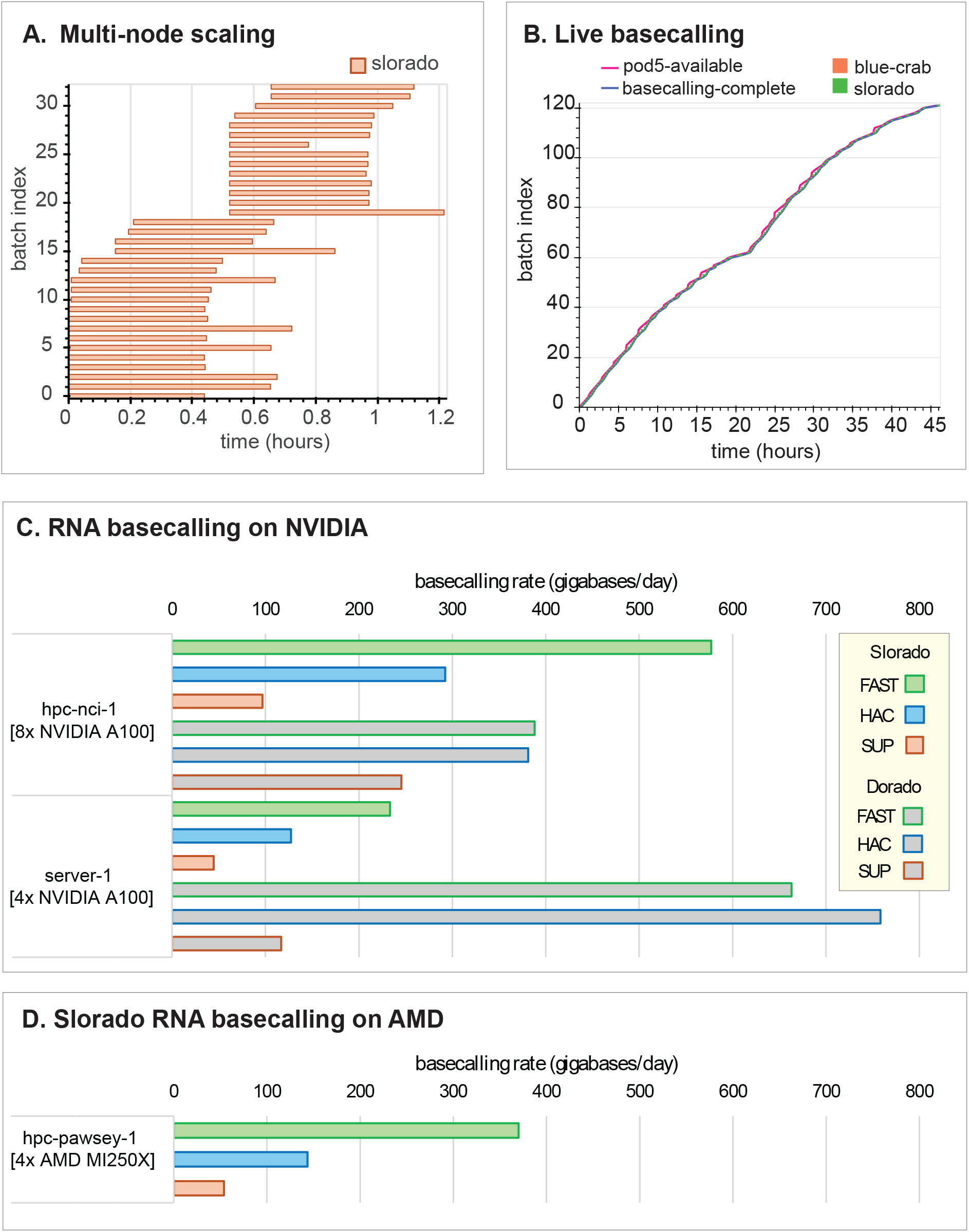
**A**. The job-start and job-finish times for the full 16.1 million read HG002-dna dataset split across 33 individual Slorado SUP v5.0.0 basecalling jobs on the Pawsey Supercomputer cluster. **B**. Real-time DNA basecalling performance of Slorado using HAC v5.0.0 over a 2-day sequencing run. The raw signal data for this run was generated by a PromethION flow cell and processed by Slorado on a single AMD RX 7900XTX GPU. **C**. RNA basecalling performance of Dorado vs. Slorado on NVIDIA GPUs. Calculated from the total time taken to basecall the complete UHRR-rna dataset with ONT’s v5.1.0 RNA basecalling models. **D**. RNA basecalling performance of Slorado on AMD GPUs. Calculated from the total time taken to basecall the complete UHRR-rna dataset with ONT’s v5.1.0 RNA basecalling models.

Accuracy metrics for all FASTQ outputs generated in these experiments are provided in Supplementary Fig. S2, Supplementary Tables S3, S4 and S5.

### 2.6. Slorado suitability for live basecalling

Another requirement for practical usage is the capacity to perform basecalling in real time, at the rate of sequencing data production on an ONT instrument. For this experiment, we simulated a full live-basecalling PromethION run over 2 days. Reads produced by the PromethION were streamed through batches in the native POD5 file format. To basecall these reads, we first translated the POD5 file into the BLOW5 file format with Blue-crab (see Supplementary Note 1), and then passed the BLOW5 file to Slorado for HAC basecalling, which was executed on a desktop (desktop-1 in Table 2) equipped with a single consumer-grade AMD RX 7900XTX GPU (retail price under 1000 USD). With this relatively limited hardware, Slorado was able to keep up with the sequencing rate of a PromethION flowcell (Fig. 3B). Across every batch in the live run, the POD5 to BLOW5 conversion accounted for only 4-7% of the total (conversion + basecalling) time (Supplementary Fig. S3). All POD5 files were completely translated to FASTQ format when the sequencing ended. This demonstrates the feasibility of performing live ONT basecalling on inexpensive hardware not currently supported using Dorado.

### 2.7. Slorado compatibility

Slorado is designed to open state-of-the-art basecalling to as many devices as possible. On top of NVIDIA and AMD platforms presented above, Slorado also supports mass-produced consumer-grade embedded GPU devices. To demonstrate Slorado’s range of support, we tested Slorado on an NVIDIA Jetson AGX Xavier (edge-1 in Table 2) and an AMD 760M integrated-GPU (edge-2 in Table 2). We were able to successfully complete FAST basecalling with Slorado on a 500K read dataset (HG002-dna-500k in Table 1) in 2.4 hours on the NVIDIA Jetson AGX Xavier and 2.4 hours on the AMD 760M integrated-GPU (see Supplementary Table S7). Dorado v1.1.1 produced an error during attempted execution on the NVIDIA Jetson AGX Xavier device, and is unsupported on AMD.

### 2.8. Slorado support for RNA basecalling

Slorado also supports RNA basecalling on the same devices supporting DNA basecalling, since they use the same underlying neural network architecture. Here, we performed RNA basecalling on a 16.4 million read dataset (UHRR-rna in Table 1) with GPU nodes hosted by NCI and Pawsey, as well as on the custom server (Fig. 3C and 3D). After basecalling, we verified the accuracy of our results (see Methods). As observed for DNA basecalling (above), Slorado and Dorado produced equivalent identity scores from RNA basecalling (Supplementary Fig. S2, Supplementary Table S6).

### 2.9. Slorado support for modification calling

Slorado additionally supports modification calling, currently for 5mC in the CpG context in DNA (ONT’s 5mCG_5hmCG@v3 model) and m6A in the DRACH motif context in RNA (m6A_DRACH@v1). To verify concordance between Slorado and Dorado for DNA 5mC, we processed the complete HG002-dna dataset (Table 1) and benchmarked the per-site methylation frequencies calculated from the output of both tools against bisulfite sequencing data (see Methods). Slorado and Dorado achieved identical Pearson correlations of 0.919 with bisulfite methylation frequencies (Supplementary Fig. S4A-B).

We then processed the complete UHRR-rna dataset (Table 1) using both Slorado and Dorado (see Methods) to assess the concordance for RNA m6A. As no orthogonal dataset was available, we compared Slorado and Dorado to each other directly, and used the run-to-run variability of Dorado itself as the baseline for how much agreement to expect, by running Dorado on two NVIDIA GPU architectures (A100 and V100) and comparing the two runs in the same way. Per-site m6A frequencies from Slorado and Dorado achieved a correlation of 0.928, a similar value for Dorado itself between two different GPU architectures (0.929), confirming that the discrepancies between Slorado and Dorado fall within the variation expected from running original Dorado on different GPUs, rather than reflecting a difference in our implementation (Supplementary Fig. S4C-D).

For both Slorado and Dorado, enabling modification detection adds a minor impact on the overall end-to-end basecalling time. We measured the end-to-end basecalling time with and without modification detection for both basecallers on the same system (see Methods). When basecalling the HG002-dna dataset, Slorado with modification detection enabled added 4% to the overall basecalling time, whereas Dorado added 1%. When basecalling the UHRR-rna dataset, Slorado with modification detection added 5% to end-to-end basecalling time, whereas Dorado added 9%.

## 3. Discussion and Future Work

The performance of ONT’s Dorado basecalling software is highly dependent on platform-specific hardware optimisations implemented in the proprietary Koi library. This effectively limits state-of-the-art ONT basecalling to be run on specific NVIDIA GPUs for which Dorado/Koi has been optimised. Such dependence on closed, hardware-specific acceleration limits portability, constrains experimentation, and prevents researchers from fully leveraging the diversity of modern computing hardware.

In this work, we have addressed this limitation by accelerating the most performance-critical component of nanopore basecalling within an accessible, open-source package, Openfish. By providing a portable implementation that supports the majority of mainstream GPU platforms, including both NVIDIA and AMD devices, Openfish removes a major barrier to experimentation and deployment. Importantly, the approach supports both RNN and attention-based basecalling models used by ONT, ensuring continued relevance as neural architectures evolve. Openfish establishes a foundation for hardware-agnostic, open-source nanopore basecalling that can track ongoing advances in model accuracy without being tightly coupled to proprietary or hardware-specific implementations.

Building on Openfish, we developed Slorado, a fully open-source end-to-end basecalling framework. Compared to Dorado, Slorado relies on far fewer dependencies (20 direct dependencies for Dorado vs 4 for Slorado; Supplementary Fig. S5) and is considerably easier to compile and deploy, enabling it to run on a wider range of devices and system configurations. This flexibility enables researchers to exploit otherwise under-utilised or more affordable hardware for nanopore signal analysis. We demonstrate that Slorado can scale across a wide spectrum of use cases, from realistic real-time ONT basecalling on a single consumer-grade GPU to parallel processing of a large ONT dataset on dozens of HPC GPU nodes for minimum turnaround time. As access to diverse compute resources continues to grow, Slorado will help lower the barrier to entry for nanopore data analysis, while enabling established users to capitalise on existing infrastructure.

In addition to Dorado, ONT provides Bonito, a Python-based basecaller primarily intended for basecalling methods development, although it also supports inference [40]. Like Dorado, Bonito relies on the proprietary Koi library that only supports NVIDIA hardware. We benchmarked Slorado against Bonito and found that Slorado achieved more than a 2*×* higher basecalling throughput with the FAST model, while delivering comparable throughput for the HAC and SUP models, with equivalent accuracy in all cases (Supplementary Fig. S6A-B).

Openfish and Slorado open the possibility of extending nanopore basecalling to additional GPU ecosystems beyond AMD and NVIDIA. As a proof of concept, we implemented a Metal backend for Openfish decoding and integrated it into Slorado to support Apple silicon, achieving performance comparable to Dorado (Supplementary Fig. S6C-D). Similarly, Openfish and Slorado provide a foundation for exploring Intel GPUs and other emerging accelerators as viable platforms for nanopore basecalling.

Besides standard basecalling, Slorado also enables modified base detection across a broader range of hardware platforms. We have implemented and validated support for the most widely used modification models for DNA (5mC) and RNA (m6A). Beyond the models currently supported, Openfish and Slorado provide a foundation for adopting future basecalling models that employ CRF-CTC decoding. For example, Openfish (requiring no additional changes) was compatible with ONT’s recently released v6 HAC models, which use a factorised LSTM-based architecture. As a proof of concept, we implemented support for the v6 HAC model in Slorado and observed a median read accuracy of 0.9709, compared with 0.9706 achieved by Dorado after basecalling the complete 16M read UHRR-rna dataset.

Beyond GPUs, the techniques described in this work apply to other parallel compute architectures. The accelerated decoding algorithm is not inherently GPU-specific and could be generalised to other platforms capable of efficient parallel execution. Platforms such as field-programmable gate arrays (FPGAs) represent a promising direction for future work, potentially offering lower latency and improved power efficiency compared to equivalently priced GPU-based systems. Exploring these platforms could further expand the applicability of open-source basecalling, particularly in embedded or edge-compute scenarios.

Currently, neural network inference exists as the dominant bottleneck in open-source nanopore basecalling pipelines. Future work will focus on sustainable optimisation strategies that balance portability, performance, and accuracy, including model compression, mixed-precision inference, and hardware-aware scheduling techniques. Continued progress in this area has the potential to substantially reduce the computational cost of nanopore sequencing and further accelerate downstream biological and clinical research.

## 4. Methods

### 4.1. Benchmarking Experiments

#### Datasets

For DNA basecalling experiments, we used a HG002 dataset sequenced on an ONT PromethION using the R10.4.1 (5kHz) chemistry (HG002-dna in Table 1), which contained ∼16.1 million reads. For use in experiments where the runtime was impractical with this full HG002-dna dataset, subsets (HG002-dna-500k and HG002-dna-20k in Table 1) were generated from the HG002-dna dataset via random selection (and reused across benchmarks). For RNA basecalling experiments, we used a Universal human RNA dataset sequenced on an ONT PromethION using the RNA004 chemistry (UHRR-rna in Table 1).

#### Computer Systems

We used several computer systems ranging from servers to edge devices for the experiments (Table 2). Server-1 is an in-house Xenon Server with NVIDIA A100 GPUs. Hpc-nci-1 and hpc-nci-2 are GPU compute nodes in the Gadi supercomputer provided by NCI, hosting NVIDIA A100 and H200 GPUs, respectively. Hpc-pawsey-1 is a GPU node in the Setonix supercomputer by Pawsey, equipped with AMD MI250X GPUs. Hpc-amd-1 is a GPU node in AMD’s AI & HPC Cluster equipped with AMD MI300X GPUs. Desktop-1 is a consumer-grade desktop hosting a single AMD RX 7900XTX. Edge-1 is a Jetson AGX Xavier edge computing device, while edge-2 is a Minisforum UM760 portable mini desktop with an integrated AMD 760M GPU.

#### Basecalling models and versions

For all DNA experiments we used ONT’s v5.0.0 models with both Slorado and Dorado. For RNA basecalling, we used v5.1.0 models. Dorado v1.1.1 and Slorado v0.4.0-beta were used for the experiments unless otherwise stated.

#### Identity Score accuracy evaluation

We evaluated the accuracy of each DNA basecalling run by using Minimap2 [41] to align basecalled reads (in FASTQ format) to the hg38 reference, and to calculate the BLAST-like identity scores for primary alignments. Similarly, we use Minimap2 to align the RNA basecalled reads to the gencode v40 transcriptome before computing the per-read identity scores from primary alignments.

#### Variant Calling accuracy evaluation

For variant calling evaluation we basecalled the complete 40*×* read depth HG002-dna-40*×* dataset (Table 1) with Slorado and Dorado using the v5.0.0 SUP model. Reads from each basecaller were aligned to hg38 with Minimap2 and sorted with Samtools. Variants were called with Clair3. Each call set was benchmarked against the GIAB HG002 v4.2.1 truthset using RTG Tools vcfeval. To assess variant-calling accuracy within homopolymer regions, we restricted the RTG Tools evaluation to the homopolymer regions defined in the GIAB genome stratifications resource (v3.6) [downloaded from https://ftp-trace.ncbi.nlm.nih.gov/ReferenceSamples/giab/release/genome-stratifications/v3.6/GRCh38@all/LowComplexity/GRCh38_AllHomopolymers_ge7bp_imperfectge11bp_slop5.bed.gz]. To focus the analysis on the homopolymer sequences themselves rather than the surrounding non-homopolymer bases, we reduced the flanking region from the 5 bases included in this file to 2 bases. Commands and software versions are provided in Supplementary Note 1.

#### Execution time measurement

The total basecalling execution time was recorded from the elapsed (wall clock) time provided by the GNU time command, available on Linux platforms. On systems without the GNU time command (hpc-pawsey-1, hpc-amd-1), the total execution time was measured using the ‘gettimeofday’ function added to the start and the end of the program source code.

#### Measuring impact of Koi

To assess how basecalling performance depends on Koi (section 2.1, Fig. 1B and 1C), we first integrated Dorado’s CPU decoder (Algorithm 1) into Slorado. This original CPU decoder did not utilise multiple CPUs efficiently, so we optimised it to fully leverage all available cores, thereby improving execution time. Then we measured the end-to-end basecalling time for the HG002-dna-20k dataset on a single NVIDIA A100 GPU on server-1 to calculate the basecalling rate (grey bars in Fig. 1B). If we had used the unmodified CPU decoder, the basecalling times would have been even more impractical. The basecalling time breakdown (Fig. 1C) was obtained using this same setup, with each major step measured using the ‘gettimeofday’ function after CPU thread synchronisation. For stages running asynchronously on the GPU, we invoked LibTorch’s ‘synchronize’ function before taking time measurements. To measure the basecalling time when closed-source Koi is used (yellow bars in Fig. 1B), we replaced the CPU decoder in Slorado with closed-source Koi binaries. To measure the basecalling time when using Openfish (section 2.2, pink bars in Fig. 1B), we replaced Koi with Openfish.

#### Single-node multi-GPU scaling analysis

To show Slorado’s ability to scale with the number of GPUs within a single node (Fig. 2C and Fig. 2D), we basecalled the HG002-dna-500k dataset (Table 1). For each iteration of the experiment, we scaled up from 1 to 8 GPUs, doubling the number of GPUs used each time (1, 2, 4, and 8 GPUs used by each basecalling model). We specified the number of GPUs to be used in Slorado using the ‘-x’ option.

#### Multi-node scaling analysis

For the experiment where we used multiple nodes available in the Pawsey supercomputer (Fig. 3A), we used Slow5tools [42] to split the complete HG002-dna (Table 1) BLOW5 file with ∼16M reads, into 33 files with ∼500K reads in each. We then submitted all 33 files simultaneously for processing with one job per file. Each job allocates a single GPU node (4*×*AMD MI250X) for SUP basecalling with Slorado. The total basecalling time was measured by recording the start time of the first job and the finishing time of the last job processed using the Linux ‘date’ command. The scripts and commands used for this are provided in Supplementary Note 1. To measure the accuracy, we first stitched the FASTQ files output by Slorado using the linux ‘cat’ command, and then we calculated the resulting identity scores using the method described above.

#### Live basecalling experiment

For the experiment where we demonstrated Slorado’s ability to perform live basecalling (Fig. 3B), we simulated a 48-hour PromethION run. This was set up so that once a POD5 file became available, blue-crab would convert it to a BLOW5 file (Supplementary Note 1) before being basecalled by Slorado. The scripts for this setup were adapted from the framework presented in [43]. We performed this experiment on desktop-1 (Table 2), which hosted a single AMD Radeon RX 7900XTX GPU. We measured the timestamps for each step using the Linux ‘date’ command. Similar to the multi-node scaling experiment above, we concatenated the output of Slorado into a single FASTQ file and calculated the basecalling accuracy using the methods described above.

#### Modification calling

For DNA modification calling, we processed the complete HG002-dna dataset (Table 1) using Slorado v0.6.0 and Dorado v2.0.0 with the SUP v5.0.0 5mCG_5hmCG@v3 model on server-1 system (Table 2). The reads were aligned to the hg38 reference using Minimap2 along with MM/ML tags. Per-site modification frequencies were calculated with Minimod [44]. The same procedure was repeated on the UHRR-rna dataset using the HAC v6.0.0 m6A_DRACH@v1 model, with the gencode v40 transcriptome used for alignment. For DNA, we compared the modification frequencies against publicly available HG002 bisulfite data from the ONT open-data AWS repository (s3://ont-open-data/gm24385_mod_2021.09/bisulphite/cpg) using the compare_methylation_ont.py and plot_methylation.R scripts available in the Slorado repository. For RNA, where orthogonal bisulfite data is not available, we compared the modification frequencies between Slorado and Dorado. As a baseline we repeated the Dorado RNA experiment on a system with NVIDIA V100 GPUs, and compared both Dorado A100 and Dorado V100 runs. All commands and software versions used for this experiment are provided in Supplementary Note 1.

### 4.2. Slorado and Openfish Implementation

Openfish implements GPU optimisations for Slorado. The GPU code in Openfish is implemented using CUDA C (for NVIDIA GPUs) and HIP C (for AMD GPUs). The CPU code in both Slorado and Openfish is written using C/C++. Slorado depends on external packages LibTorch, Slow5lib and Openfish. LibTorch is an open-source neural-network library used in Slorado for hosting ONT’s basecalling models and performing tensor operations. Slow5lib is an open-source library for reading the input data in the SLOW5 file format. CPU multithreading in Slorado is implemented using POSIX threads with a fork-join model. Slorado constructs the neural network from weights and configurations available in basecalling models released by ONT. The input SLOW5 files are then basecalled and output in the FASTQ file format.

### 4.3. Optimisation of decoding using GPU

We begin by describing the original CPU-based implementation of the decoding algorithm in Dorado, which, to the best of our knowledge, has not been formally documented or explained elsewhere. We then present our GPU-optimised adaptation of this algorithm, as implemented in Openfish.

#### 4.3.1. Original CPU Decoder

Each major step of the original CPU decoder in Dorado is outlined in Algorithm 1. Given the input matrix **scores** ∈ ℝ^*T ×N ×C*^ (generated from the inference step), where *T* is the number of time steps in our sequence, *N* is the batch size, and *C* is the size of the transitional state space, the algorithm produces a predicted DNA sequence and the Phred quality score (probability of error) of each base in the sequence (Fig. 4A). This is represented as **seqs** ∈ {A,T,C,G} ^*N ×T*^ and **qscore** ∈ ℝ^*N ×T*^, respectively. Here, each chunk *n* in batch [0..*N* ) is processed independently, limited by the number of parallel threads on the CPU. Additionally, the number of nucleotide bases is denoted as *nbase* = 4, and the number of possible states as *nstate* = *nbase*^*state*_*length*^ (where *state*_*length* is provided by the basecalling model).

**Fig. 4:**
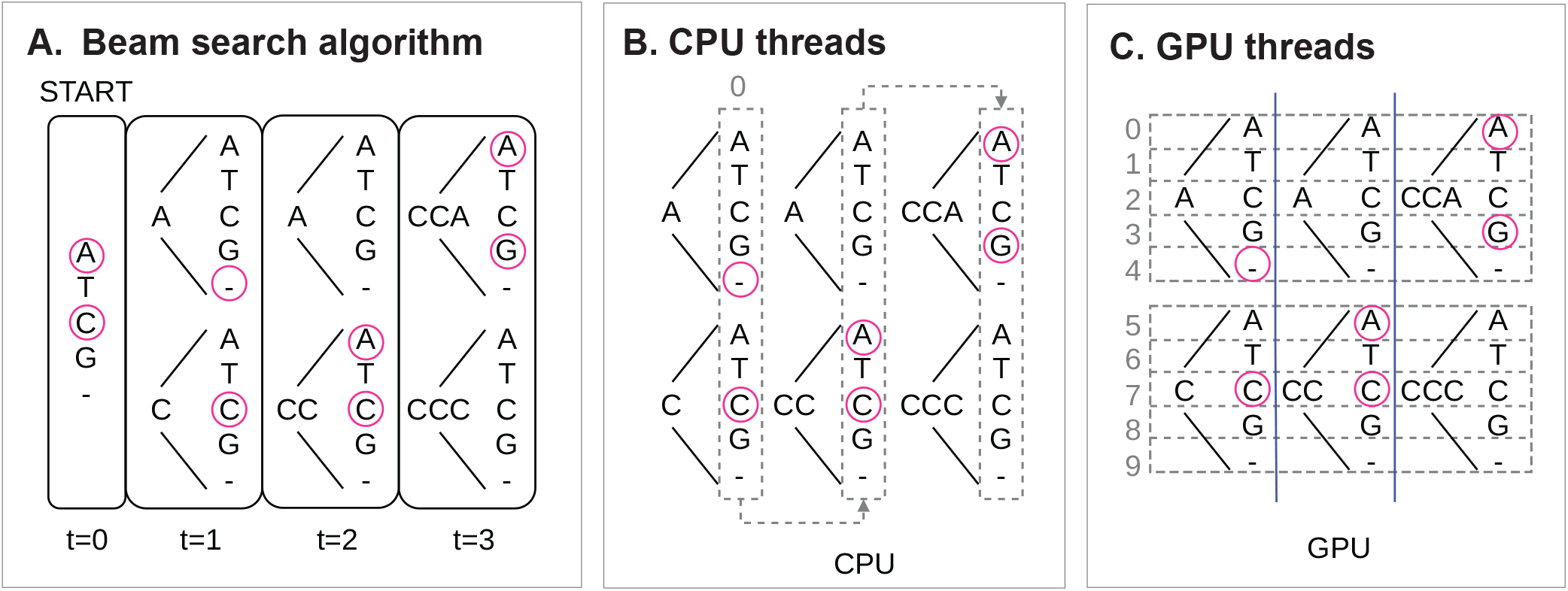
**A**. A simplified example demonstrating the execution of a beam search algorithm with a beam width=2 and a state length=1. At each time step (*t*), the best-scoring state transitions are selected to build the top paths (beams) for the final DNA sequence. In this example, the top 2 paths are built equal to the beam width. **B**. The thread execution model of the original CPU implementation of beam search. Each beam candidate (reachable state) is scored sequentially on a single thread. **C**. A simplified version of the GPU thread execution model implemented in Openfish. Multiple threads are used to score beam candidates in parallel (synchronised at the blue line). Each chunk of the neural network output is processed in a single thread block on the GPU. At the end of each time step, the top n scoring beams are found to keep in the search.

##### Algorithm 1 CPU_decode

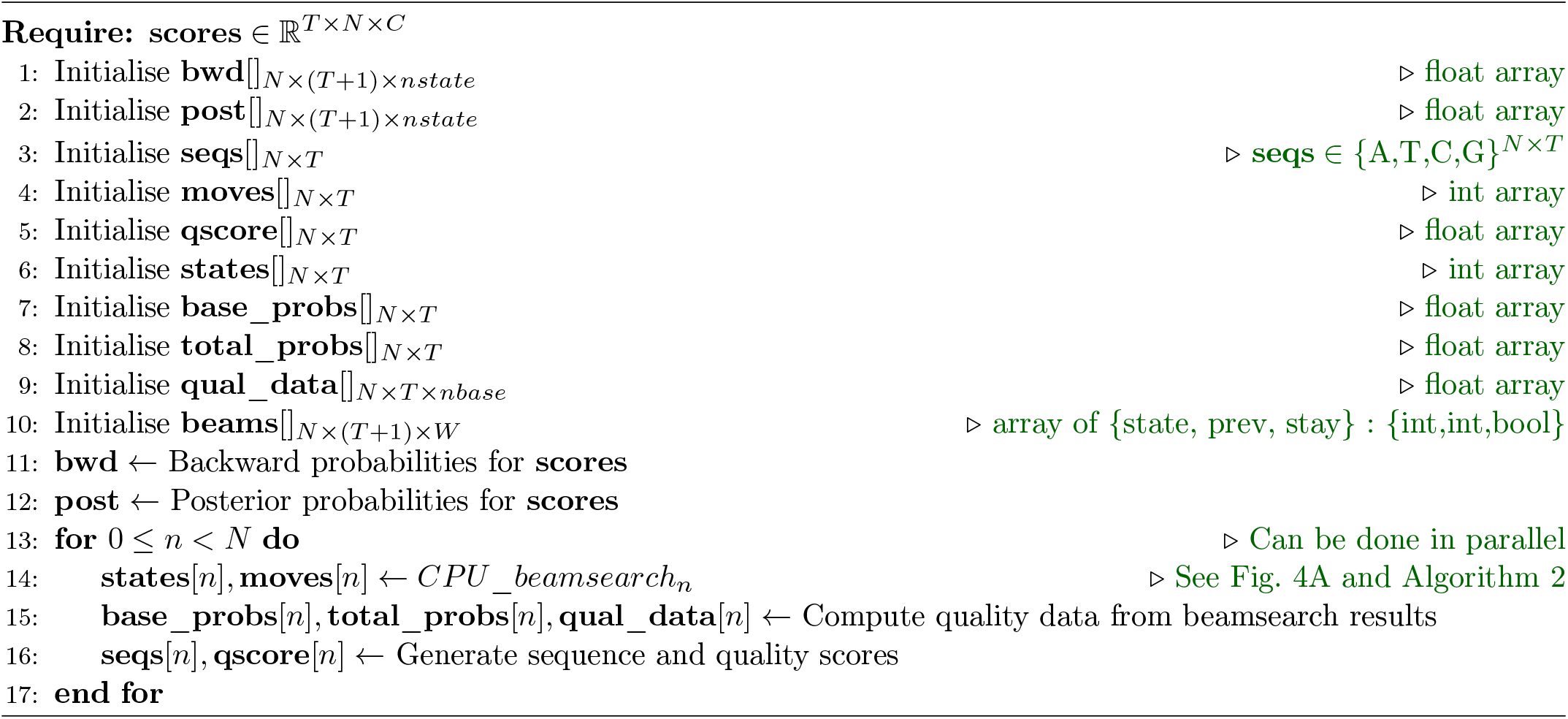

The beam search algorithm shown in Fig. 4A is a crucial step in decoding; it handles the task of finding the most likely sequence of DNA bases (A, T, C, G) that can be mapped back to the signal data scored by our neural network. The nanopore signal is split into discrete sections and scored along the temporal dimension *T* . At a high level, the beam search algorithm begins by finding the top-scoring *W* states at the beginning of the chunk, where *W* is the maximum beam width (typically 32). Then, for each time step, the algorithm searches for the next best *W* states that can be reached from the previous beam front.

By keeping track of the cumulative scores of each sequence of states generated, the result is a top-scoring set of paths that can be considered for the final nucleobase sequence. Once this is complete, each step of the best-scoring path is revisited to construct the final DNA/RNA base sequence.

In Dorado’s open-source code, beam search is implemented only for the CPU. Because of this limitation, the neural network scores required for decoding must first be moved from GPU memory to CPU memory. As shown in Fig. 1C, a significant amount of memory is copied that bottlenecks the performance of the entire basecalling pipeline. Additionally, because of the volume of data that needs to be processed, beam search takes a considerable amount of time on the CPU and is a non-trivial task to port to GPU architectures.

#### 4.3.2. Openfish GPU Decoder

Modern GPUs contain thousands of computing cores for high-throughput parallel processing. The CPU beam search algorithm outlined in Fig. 4B (Algorithm 2) is not inherently suitable for this type of processing. The code executes synchronously (limited to a single thread of execution) and thus cannot sufficiently utilise the thousands of threads available to the GPU. Simply launching a thread for each chunk from [0..*N* ) in parallel on the GPU is not a sufficient optimisation, as: (1) The number of GPU threads (thousands) available for execution is typically higher than the batch size *N* (hundreds), (2) In the current CPU algorithm, each thread of execution will require its own resources and access to global memory, leading to inefficient memory allocation and expensive memory access patterns that slow down performance. Furthermore, the CPU implementation requires dynamically allocated arrays in RAM for each beam search thread. Dynamically allocating and accessing global memory in GPUs is an expensive task and should be avoided when possible.

We outline how Openfish performs beam search on the neural net scores (data generated by the inference step) without transferring data to host memory. To avoid dynamic memory allocation on the GPU, the heuristics outlined in Algorithm 3 and 4 are used to allocate and re-use statically sized memory configured from the start of the basecalling program. To address the inherent synchronous nature of the original CPU beam search algorithm (Algorithm 2), the GPU version of beam search is configured to add and score possible state transitions in parallel at each time step. This way, computational resources on the GPU can be utilised more effectively.

When searching for the best approximate DNA sequence, the number of possible paths under consideration is kept within the maximum beam width *W* . Thus, the maximum number of beam candidates we can add to each path is equal to *I* = *W ×* (*nbase* + 1) (1 is added because there is the possibility that a new base is not encountered at every time step). The number of GPU threads used for each chunk is *K* = *W × nbase*.

The scores reside in GPU Global Memory (GM). GPU threads of the same block operate on the same GPU Shared Memory (SM), which provides significantly faster read and write operations than Global Memory.

##### Algorithm 2 CPU _beamsearch

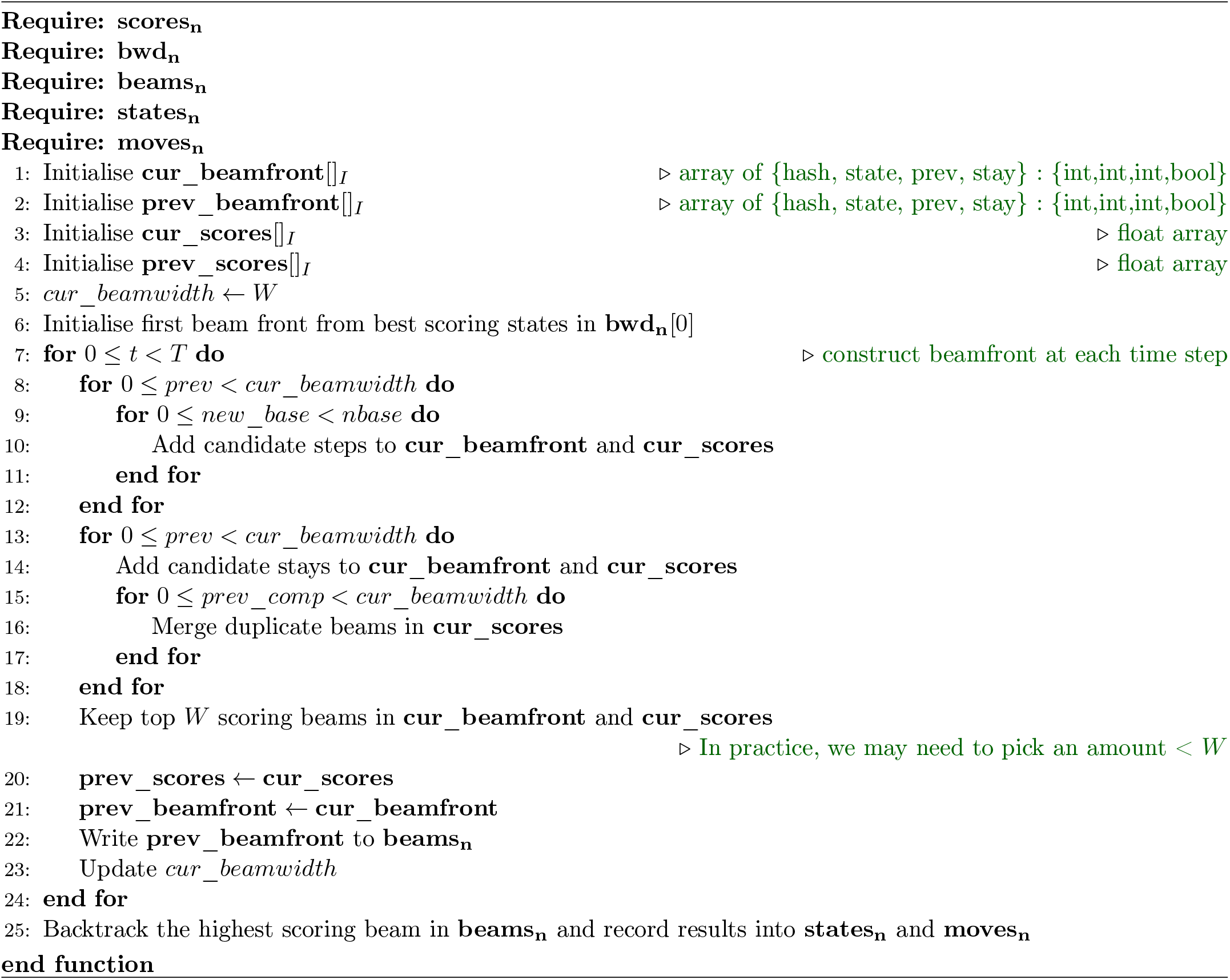

##### Algorithm 3 GPU_decode

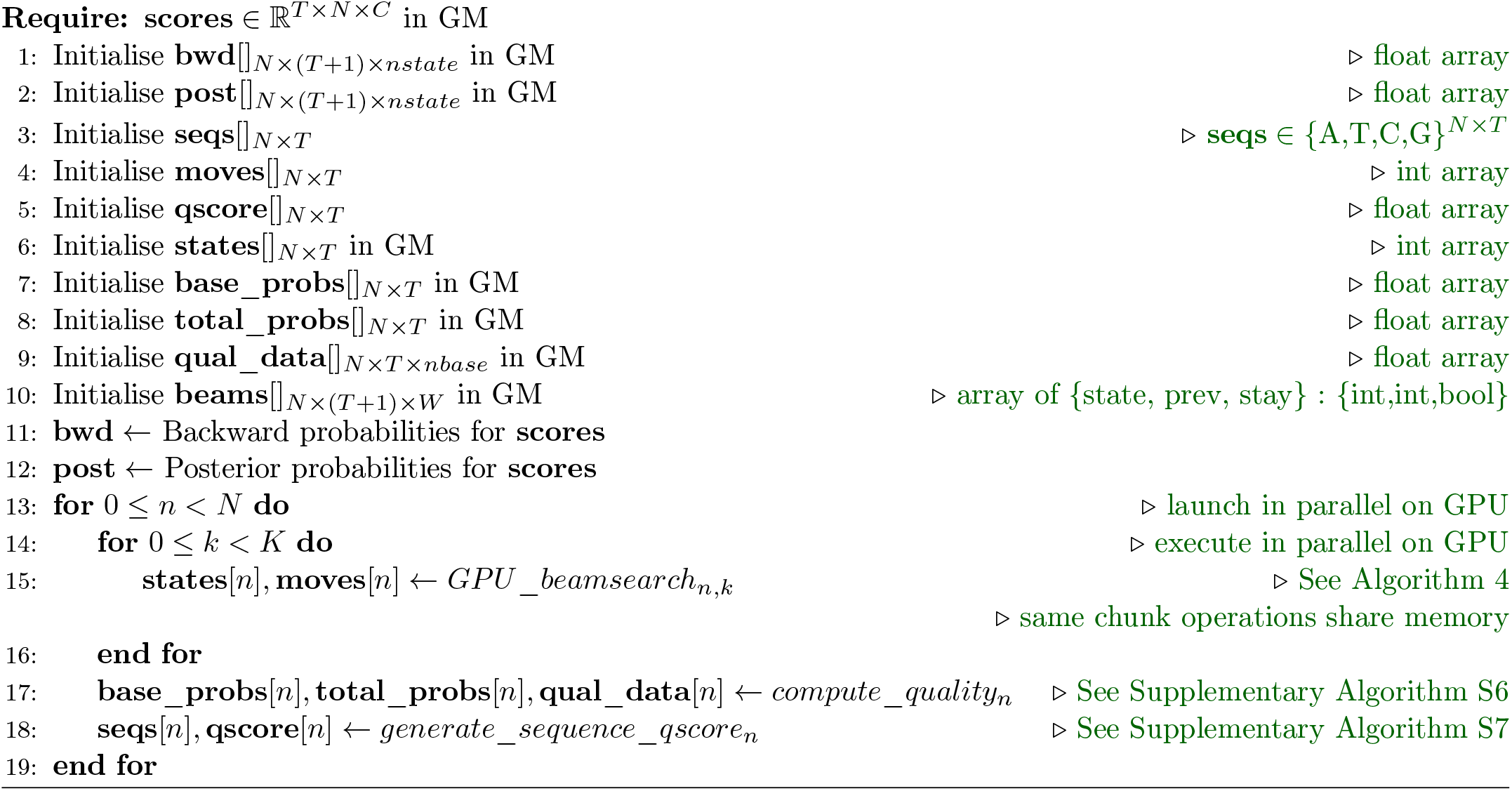

##### Algorithm 4 GPU _beamsearch

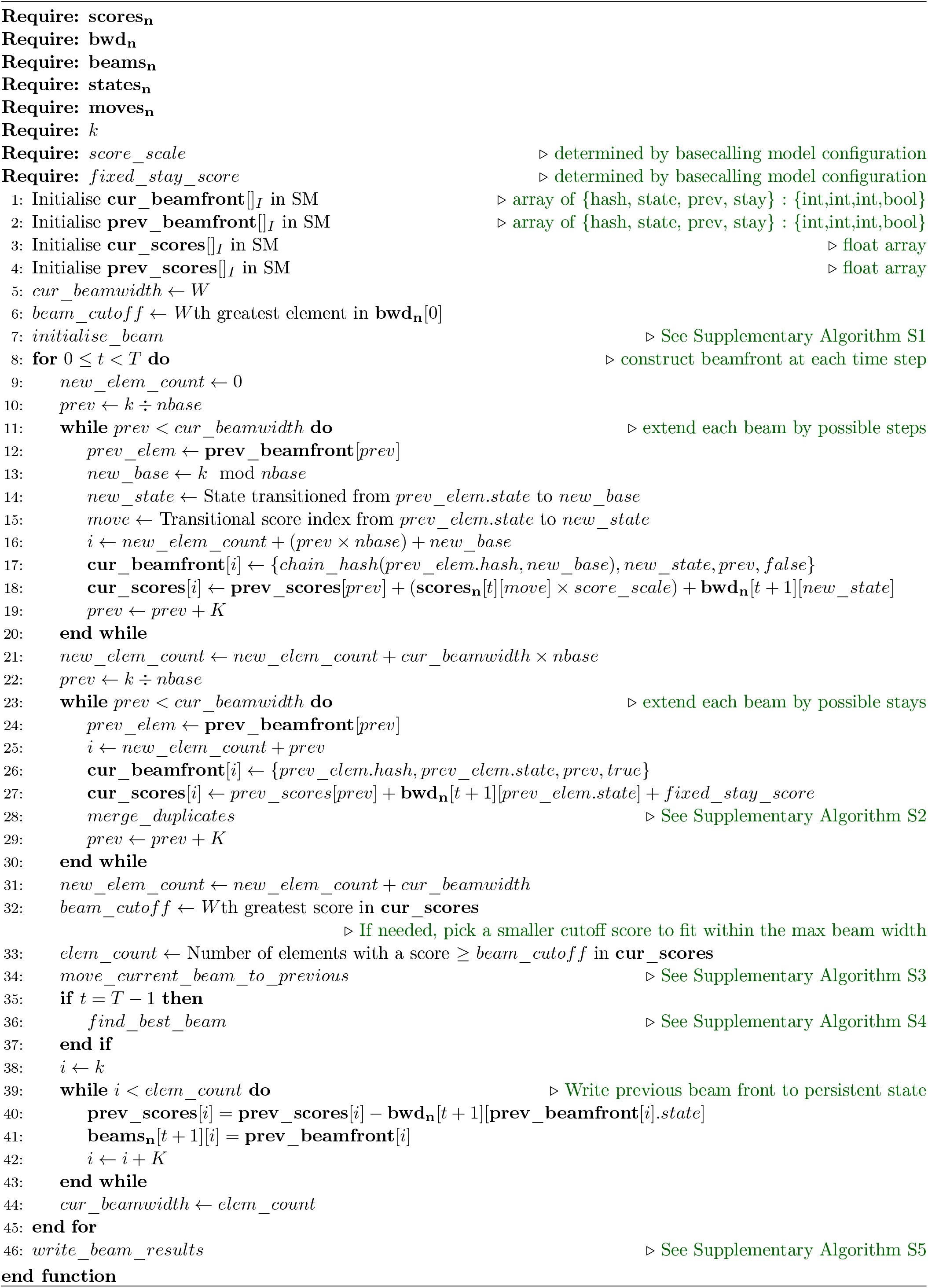

## 6. Author contributions statement

B.W., H.S., I.W.D. and H.G. conceived the project. B.W. and H.G. developed the Openfish and Slorado software. G.S., H.J. and K.D. contributed to the software design. B.W., K.L. and H.S. performed benchmarking experiments with contributions from all co-authors. B.W., I.W.D. and H.G. prepared the figures and manuscript, with contributions from G.S., H.J. and K.D. All authors read and approved the final manuscript.

## 7. Acknowledgments

We acknowledge the support from: James Ferguson (Garvan Institute of Medical Research) for building the custom computer; Hassaan Saadat (UNSW Sydney) for establishing a connection with AMD; and, Shiraun K Jacob (UNSW Sydney) for software licensing. We acknowledge the funding from the Australian Research Council DECRA Fellowship DE230100178 (to H.G.). This work was supported in part by AMD through funding as an unrestricted gift; Heterogeneous Accelerated Compute Clusters (HACC) program; and AI & HPC Cluster. Resources from NCI Australia and Pawsey Supercomputing Research Centre were used for this work. Access to Pawsey was provisioned by the Australian BioCommons Leadership Share (ABLeS) program [45]. The views expressed herein are those of the authors and are not necessarily those of the funding bodies.

## 8. Data and code availability

The source code for Openfish and Slorado is available as open-source at https://github.com/BonsonW/slorado, respectively. The complete HG002-dna, HG002-dna-40*×* and UHRR-rna datasets used for the experiments are publicly available from the European Nucleotide Archive under the run accession numbers ERR12997167, ERR14751980 and ERR12997170, respectively.

## Supplementary materials

**Fig. S1:**
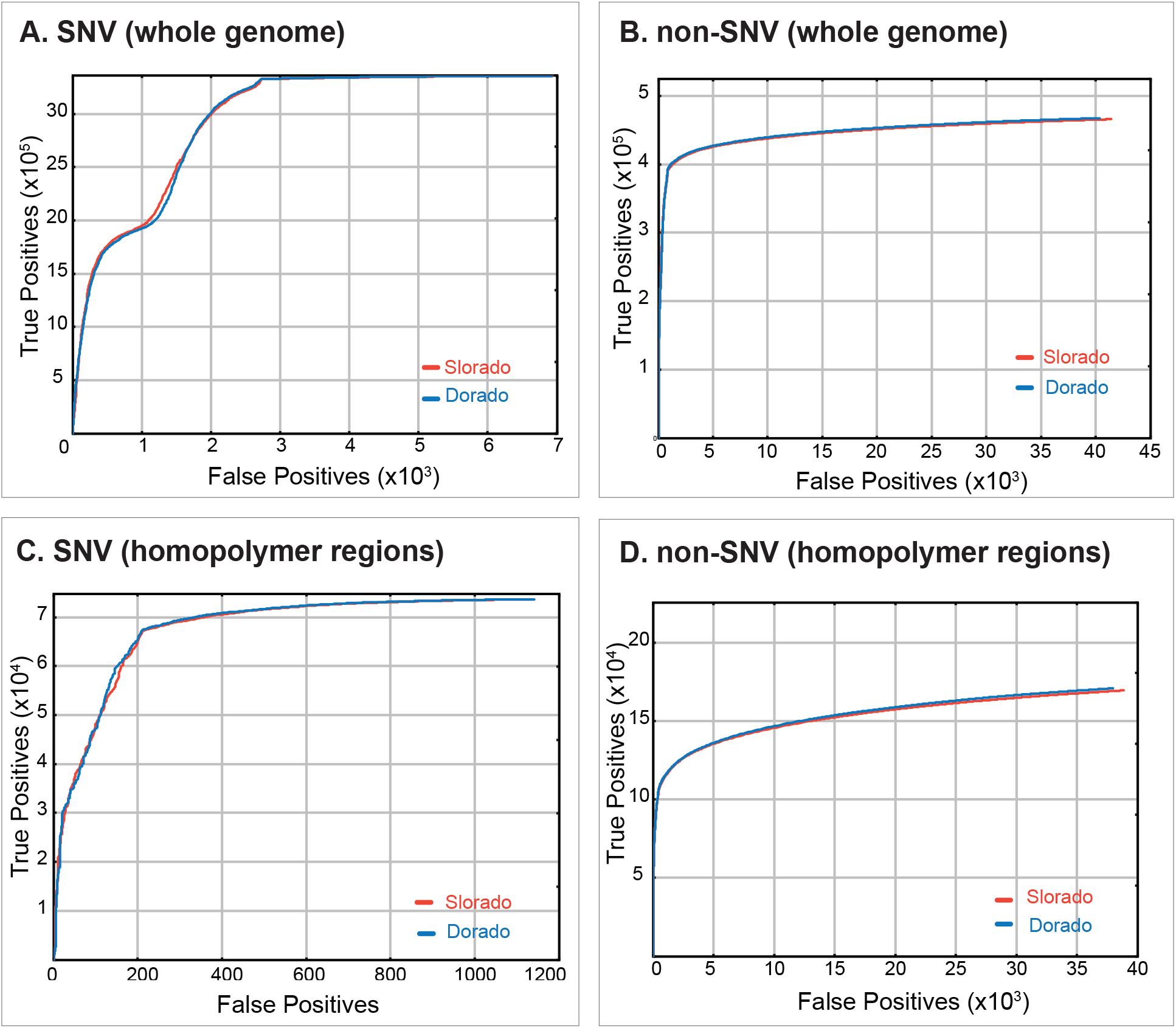
ROC curves for small variants called with Clair3 from Slorado and Dorado v5.0.0 SUP basecalls of the complete 7.2 million read HG002-dna-40*×* dataset, benchmarked against the GIAB HG002 v4.2.1 truth set. **A**. SNV, genome-wide. **B**. Non-SNV (short indels), genome-wide. **C**. SNV, homopolymer regions. **D**. Non-SNV, homopolymer regions.

**Fig. S2:**
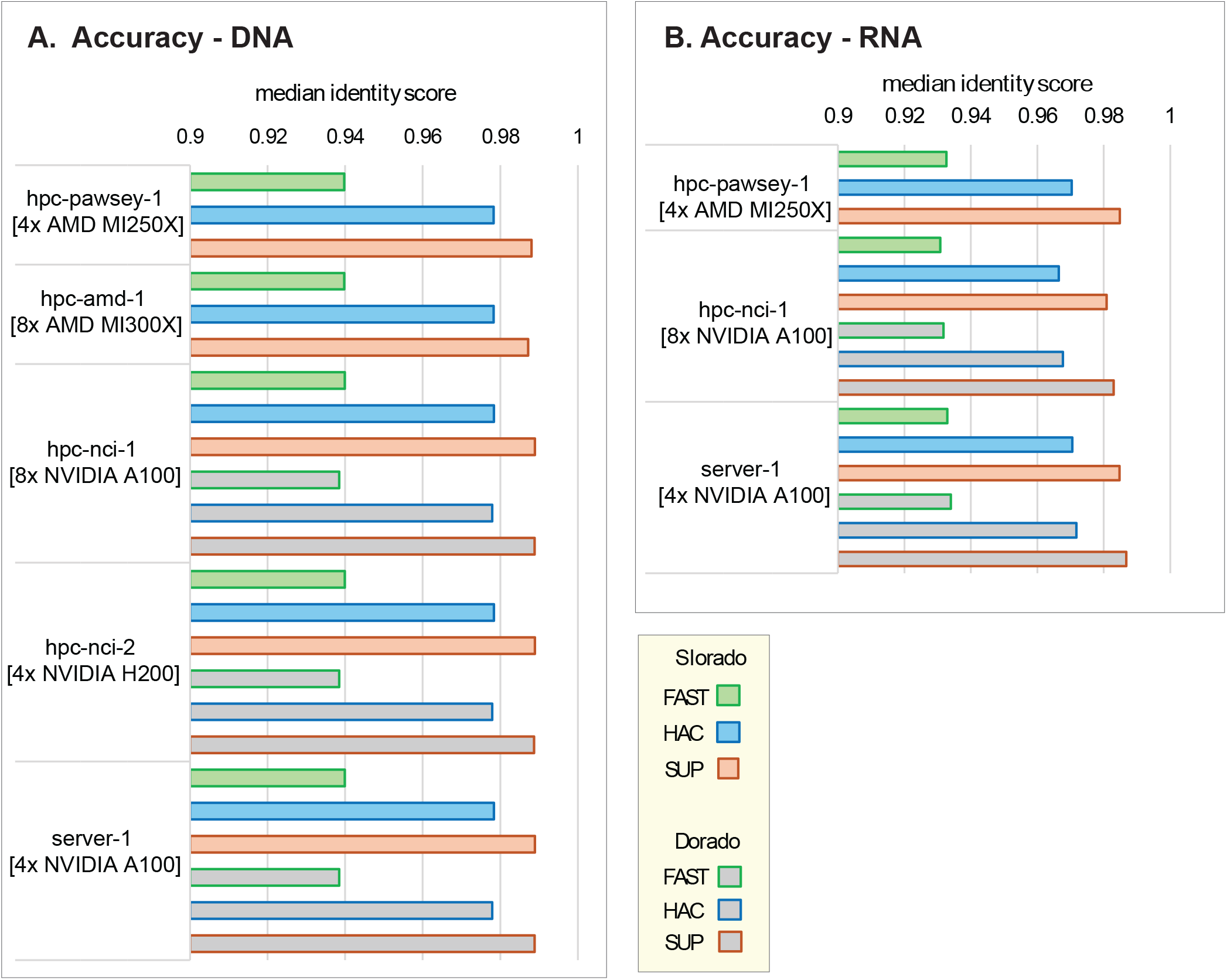
**A**. DNA basecalling accuracy comparison between Slorado v0.4.0-beta and Dorado v1.1.1 for v5.0.0 DNA basecalling models. Identity scores were collected from basecalling the complete 16.1 million read HG002-dna dataset. **B**. RNA basecalling accuracy comparison between Slorado v0.4.0 and Dorado v1.1.1 for v5.1.0 RNA basecalling models. Identity scores were collected from basecalling the complete 16.4 million read UHRR-rna dataset.

**Fig. S3:**
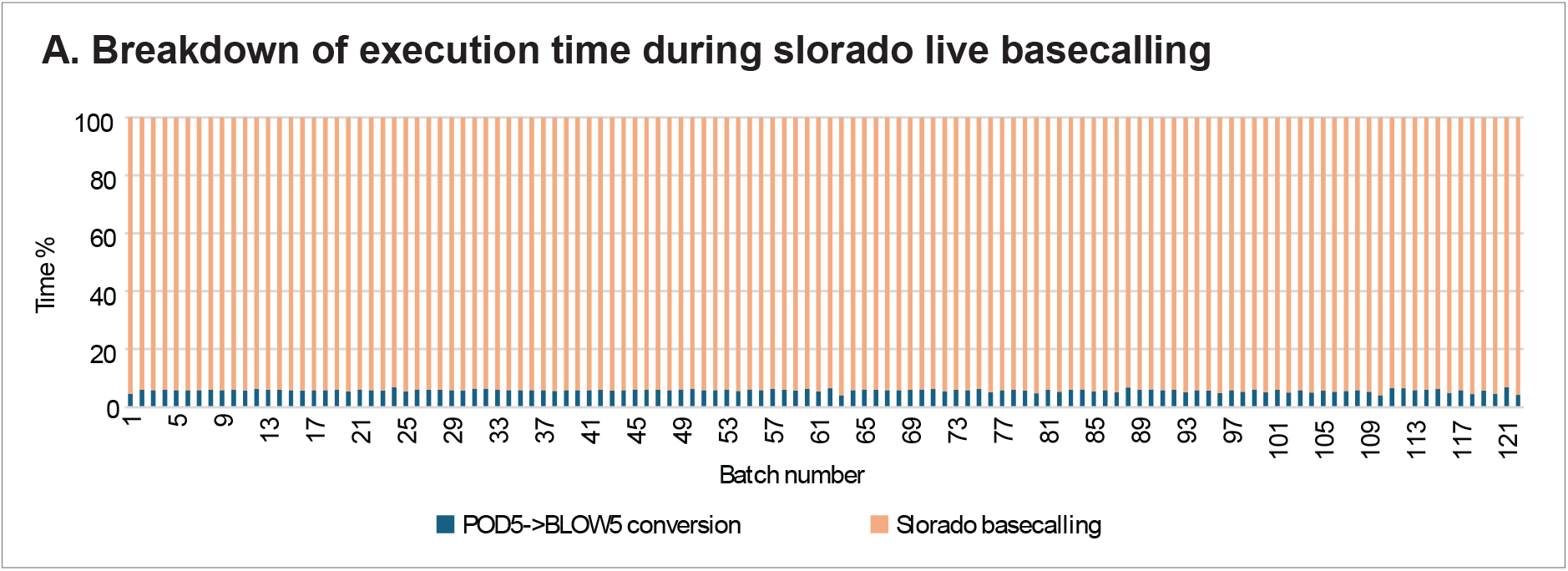
**A**. Breakdown of the time spent converting POD5 files to BLOW5 before basecalling for each batch produced in the live basecalling experiment; performed on the desktop-1 system with a single AMD RX 7900XTX and 16 CPU threads. Reads were converted to BLOW5 with Blue-crab and basecalled with Slorado using the HAC v5.0.0 DNA basecalling model.

**Fig. S4:**
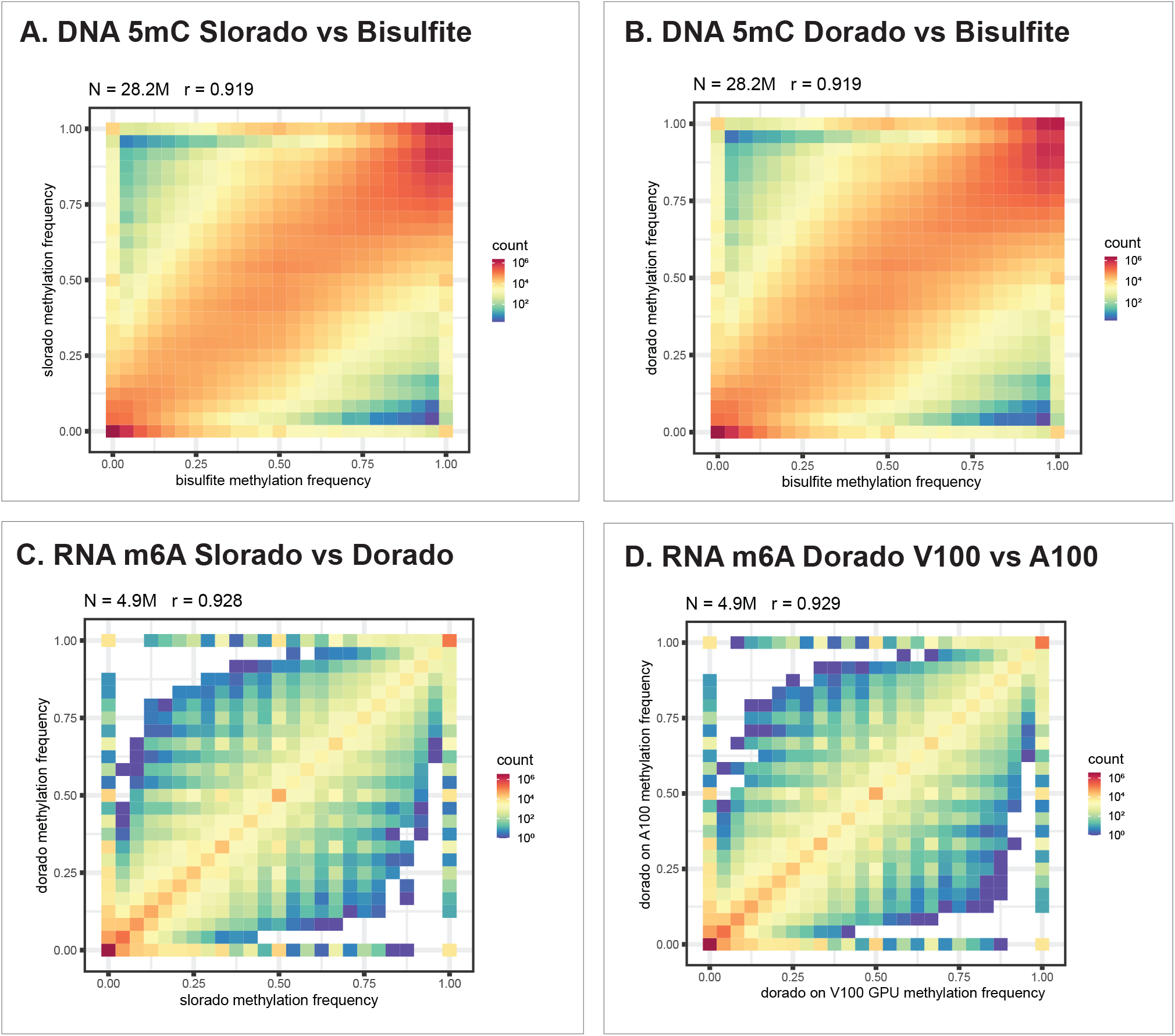
Per-site modification frequencies compared between Slorado and Dorado. DNA was basecalled from the complete 16.1M read HG002-dna dataset with the v5.0.0 SUP model and the 5mCG_5hmCG@v3 modification model; RNA was basecalled from the complete 16.4M read UHRR-rna dataset with the v6.0.0 HAC model and the m6A_DRACH@v1 modification model. **A**. DNA 5mC per-site methylation frequencies, Slorado against bisulfite sequencing of the same sample. **B**. DNA 5mC per-site methylation frequencies, Dorado against the same bisulfite data. **C**. RNA m6A per-site modification frequencies, Slorado against Dorado, both run on NVIDIA A100 GPU. **D**. RNA m6A per-site modification frequencies, Dorado run on NVIDIA A100 against the same version of Dorado run on NVIDIA V100.

**Fig. S5:**
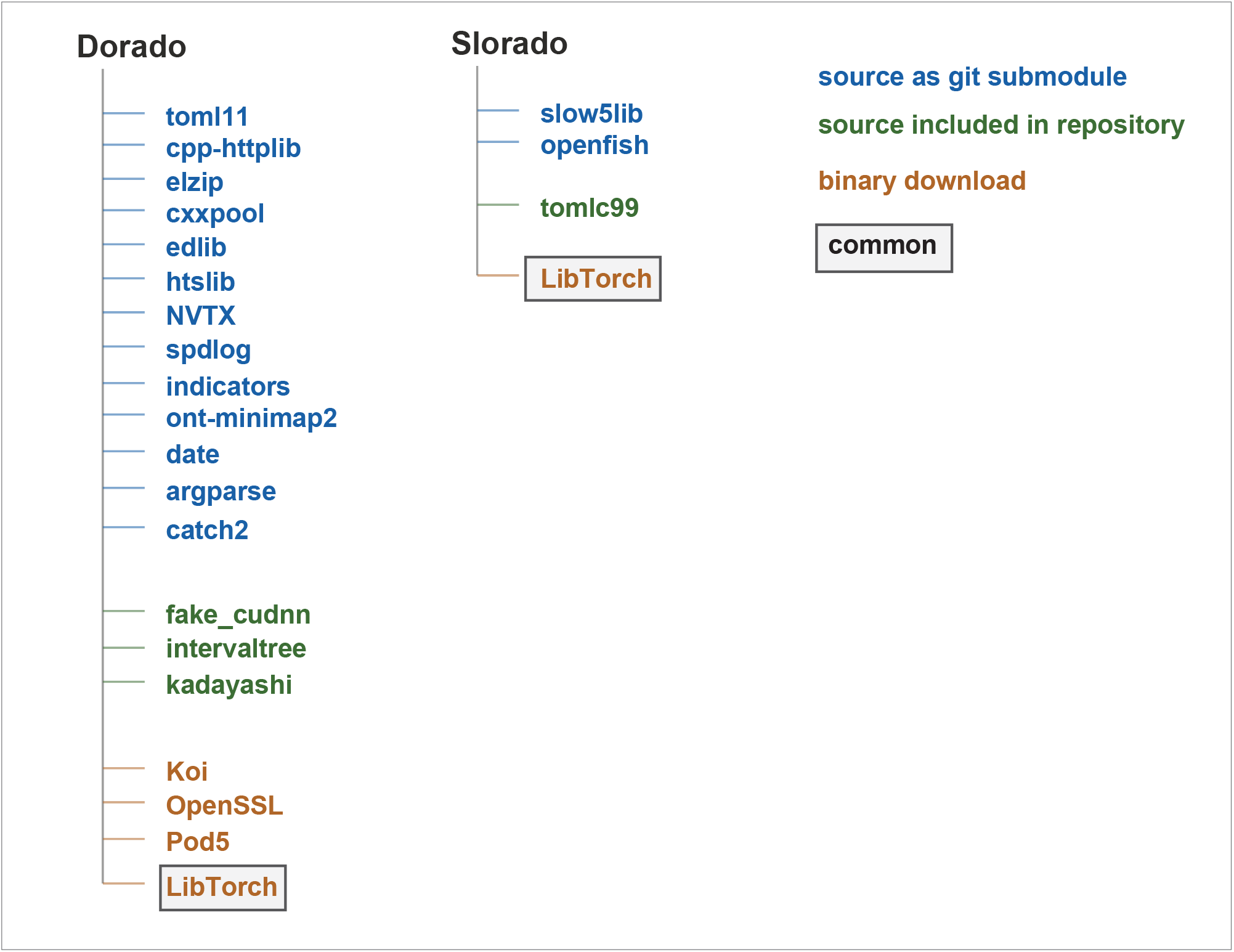
Dependencies of Dorado v1.1.1 and Slorado v0.4.0-beta. Source dependencies are compiled from code present in the working tree (depth of 1), and are split into git submodules and vendored copies included in the git repository. Binary dependencies are prebuilt and are never compiled from in-tree source.

**Fig. S6:**
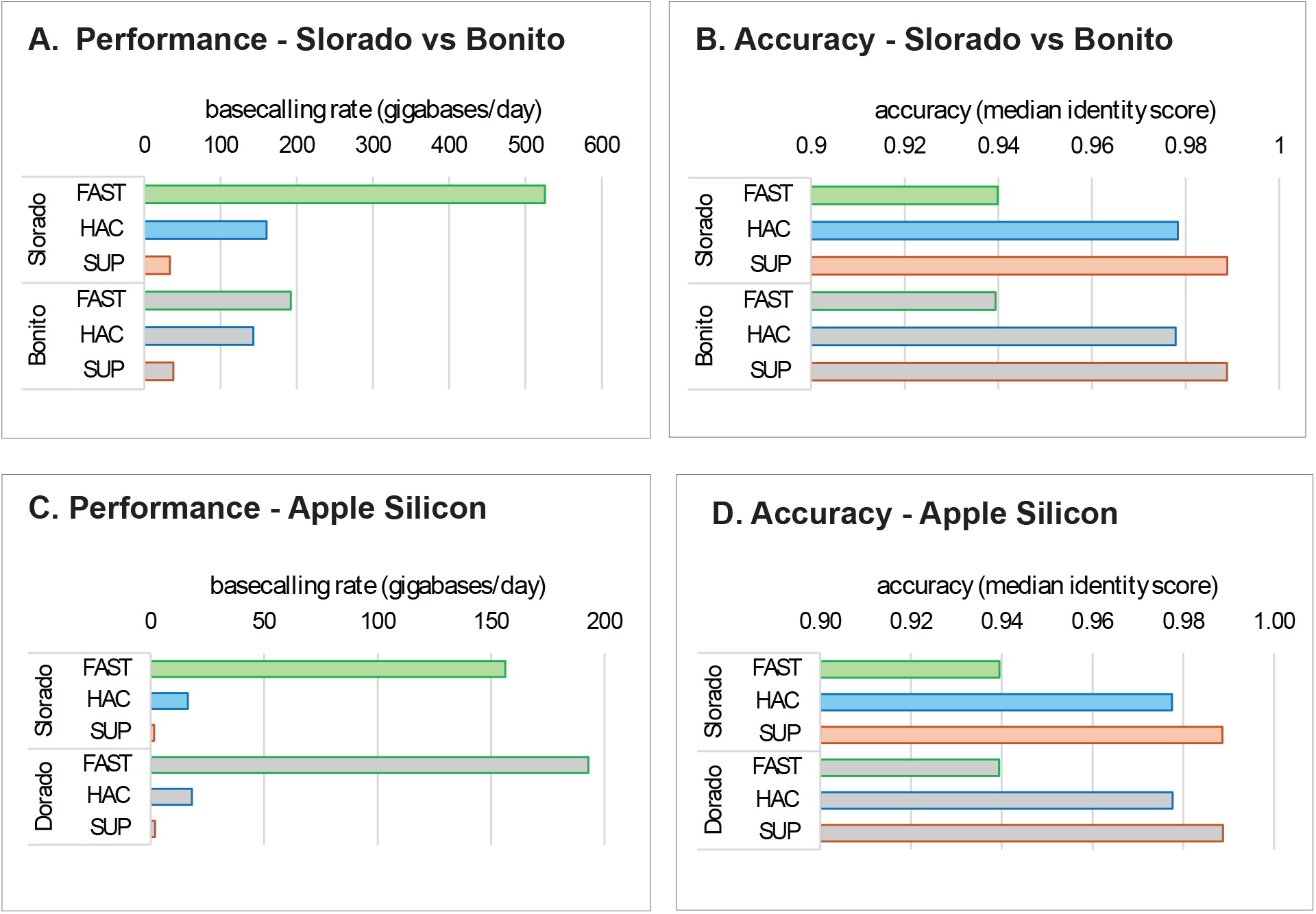
**(A)** Performance comparison and **(B)** accuracy comparison between Slorado v0.4.0-beta and Bonito v1.1.0 for ONT’s v5.0.0 DNA basecalling models. Both Slorado and Bonito were benchmarked on a single GPU on the server-1 system to ensure a fair comparison, as Bonito does not support parallel execution across multiple GPUs. The 500k read HG002-dna-500k dataset was used. **(C)** Performance comparison and **(D)** Accuracy comparison between Slorado and Dorado v1.1.1 for ONT’s v5.0.0 DNA basecalling models on an Apple M4 Pro Mac instance on Amazon AWS (14-core CPU, 20-core GPU, 48 GB unified memory, APFS Apple SSD, Sequoia 15.7.7 OS, LibTorch 2.4.0 and Metal 3). Identity scores were collected from basecalling the 20,000-read HG002-dna-20k dataset. The “metal” branch (commit db59b5b8a0149eb1694617ac3dcbd323487837d4) in the Slorado GitHub repository was used for all experiments on Apple Silicon.

**Table S1.** Basecalling time for different configurations of Koi and Openfish. Executed on server-1 using only one NVIDIA A100 GPU. Basecalling v5.0.0 models were used. HG002-dna-20k was used as the dataset.

| model | configuration | basecalling time (seconds) |
| --- | --- | --- |
| FAST | without Koi | 444 |
|  | with Koi | 28 |
|  | with Openfish | 32 |
| HAC | without Koi | 1985 |
|  | with Koi | 93 |
|  | with Openfish | 97 |
| SUP | without Koi | 6143 |
|  | with Koi | 434 |
|  | with Openfish | 447 |

**Table S2.** Breakdown of the basecalling time when executing the configuration without Koi. All values are in seconds. Executed on server-1 using only one NVIDIA A100 GPU. Basecalling v5.0.0 models were used. HG002-dna-20k was used as the dataset.

| model | inference | CPU-GPU data transfer | decode | other |
| --- | --- | --- | --- | --- |
| FAST | 16 | 211 | 159 | 58 |
| HAC | 75 | 1187 | 539 | 183 |
| SUP | 406 | 3146 | 1946 | 645 |

**Table S3.** Total basecalling time and accuracy of basecalling results from Slorado and Dorado for the HG002-dna dataset.

| system | software | model | time (hours) | accuracy<br>(median<br>identity score) |
| --- | --- | --- | --- | --- |
| hpc-pawsey-1 [4× AMD MI250X] | Slorado | FAST | 2.3 | 0.939759 |
|  |  | HAC | 5.4 | 0.978297 |
|  |  | SUP | 15.4 | 0.988082 |
| hpc-amd-1 [8× AMD MI300X] | Slorado | FAST | 0.8 | 0.939759 |
|  |  | HAC | 2.3 | 0.978300 |
|  |  | SUP | 3.7 | 0.987220 |
| hpc-nci-1 [8× NVIDIA A100] | Slorado | FAST | 1.5 | 0.939902 |
|  |  | HAC | 2.7 | 0.978336 |
|  |  | SUP | 7.6 | 0.988923 |
|  | Dorado | FAST | 2.4 | 0.938452 |
|  |  | HAC | 2.3 | 0.977957 |
|  |  | SUP | 2.7 | 0.988889 |
| hpc-nci-2 [4 × NVIDIA H200] | Slorado | FAST | 0.8 | 0.939903 |
|  |  | HAC | 1.6 | 0.978337 |
|  |  | SUP | 7.4 | 0.988920 |
|  | Dorado | FAST | 1.9 | 0.938452 |
|  |  | HAC | 1.8 | 0.977957 |
|  |  | SUP | 2.8 | 0.988701 |
| server-1 [4 × NVIDIA A100] | Slorado | FAST | 1.8 | 0.939902 |
|  |  | HAC | 3.8 | 0.978336 |
|  |  | SUP | 15.6 | 0.988923 |
|  | Dorado | FAST | 0.7 | 0.938452 |
|  |  | HAC | 0.8 | 0.977957 |
|  |  | SUP | 5.7 | 0.988889 |

**Table S4.** Total basecalling time and accuracy when using Slorado with different numbers of AMD GPUs. The experiment was conducted using the HG002-dna-500k dataset on the hpc-amd-1 system with 8*×* AMD MI300X GPUs.

| model | no. GPU | time (minutes) | accuracy<br>(median<br>identity score) |
| --- | --- | --- | --- |
| FAST | 1 | 13.0 | 0.939708 |
|  | 2 | 5.7 | 0.939708 |
|  | 4 | 3.3 | 0.939708 |
|  | 8 | 2.0 | 0.939708 |
| HAC | 1 | 37.9 | 0.978283 |
|  | 2 | 18.2 | 0.978283 |
|  | 4 | 9.7 | 0.978283 |
|  | 8 | 5.3 | 0.978283 |
| SUP | 1 | 59.7 | 0.987207 |
|  | 2 | 30.2 | 0.987207 |
|  | 4 | 15.7 | 0.987207 |
|  | 8 | 8.4 | 0.987207 |

**Table S5.** Total basecalling time and accuracy when using Slorado with different numbers of NVIDIA GPUs. The experiment was conducted using the HG002-dna-500k dataset on the hpc-nci-1 system with 8*×* NVIDIA A100 GPUs.

| model | no. GPU | time (minutes) | accuracy<br>(median<br>identity score) |
| --- | --- | --- | --- |
| FAST | 1 | 8.7 | 0.939834 |
|  | 2 | 4.3 | 0.939834 |
|  | 4 | 2.9 | 0.939834 |
|  | 8 | 2.0 | 0.939834 |
| HAC | 1 | 25.1 | 0.978309 |
|  | 2 | 13.2 | 0.978309 |
|  | 4 | 7.3 | 0.978309 |
|  | 8 | 4.4 | 0.978309 |
| SUP | 1 | 116.6 | 0.988920 |
|  | 2 | 58.9 | 0.988920 |
|  | 4 | 30.7 | 0.988920 |
|  | 8 | 16.4 | 0.988920 |

**Table S6.** Total basecalling time and accuracy of basecalling results from Slorado and Dorado for the UHRR-rna dataset.

| system | software | model | time (hours) | accuracy<br>(median<br>identity score) |
| --- | --- | --- | --- | --- |
| hpc-pawsey-1 [4× AMD MI250X] | Slorado | FAST | 1.4 | 0.932609 |
|  |  | HAC | 3.5 | 0.970402 |
|  |  | SUP | 9.2 | 0.984848 |
| hpc-nci-1 [8× NVIDIA A100] | Slorado | FAST | 0.9 | 0.930758 |
|  |  | HAC | 1.7 | 0.966516 |
|  |  | SUP | 5.2 | 0.980895 |
|  | Dorado | FAST | 1.3 | 0.931746 |
|  |  | HAC | 1.3 | 0.967692 |
|  |  | SUP | 2.0 | 0.982955 |
| server-1 [4× NVIDIA A100] | Slorado | FAST | 2.1 | 0.932897 |
|  |  | HAC | 3.9 | 0.970536 |
|  |  | SUP | 11.2 | 0.984772 |
|  | Dorado | FAST | 0.8 | 0.934019 |
|  |  | HAC | 0.7 | 0.971751 |
|  |  | SUP | 4.3 | 0.986784 |

**Table S7.** Total basecalling time and accuracy when using Slorado on edge GPU devices. The experiment was conducted using the HG002-dna-500k dataset.

| system | model | time (minutes) | accuracy<br>(median<br>identity score) |
| --- | --- | --- | --- |
| edge-1 [1 × Jetson AGX Xavier] | FAST | 144.4 | 0.939837 |
| edge-2 [1 × AMD 760M] | FAST | 141.2 | 0.939716 |

### Supplementary Note 1 - Commands and versions

#### Basecalling Options

Supplementary Note 1 - Commands and versions Basecalling Options
| system | model | no. threads | max bytes (-B) | no. max reads (-K) | chunk size (-c) | GPU batch size (-C) |
| --- | --- | --- | --- | --- | --- | --- |
| server-1 | FAST | 64 | 1G | 20000 | 10000 | 2000 |
|  | HAC | 64 | 1G | 20000 | 10000 | 2000 |
|  | SUP | 64 | 1G | 20000 | 10000 | 1000 |
| hpc-pawsey-1 | FAST | 64 | 1G | 20000 | 10000 | 2048 |
|  | HAC | 64 | 1G | 20000 | 10000 | 512 |
|  | SUP | 64 | 1G | 20000 | 10000 | 1024 |
| hpc-amd-1 | FAST | 128 | 8G | 160000 | 10000 | 2000 |
|  | HAC | 128 | 8G | 160000 | 10000 | 500 |
|  | SUP | 128 | 8G | 160000 | 10000 | 4000 |
| hpc-nci-1 | FAST | 48 | 8G | 160000 | 10000 | 4000 |
|  | HAC | 48 | 8G | 160000 | 10000 | 4000 |
|  | SUP | 48 | 8G | 160000 | 10000 | 1400 |
| hpc-nci-2 | FAST | 48 | 8G | 160000 | 10000 | 4000 |
|  | HAC | 48 | 8G | 160000 | 10000 | 4000 |
|  | SUP | 48 | 8G | 160000 | 10000 | 3000 |
| desktop-1 | HAC | 16 | 1G | 20000 | 10000 | 500 |
| edge-1 | FAST | 8 | 500M | 2000 | 10000 | 200 |
| edge-2 | FAST | 8 | 500M | 2000 | 10000 | 500 |

#### Slorado Basecalling

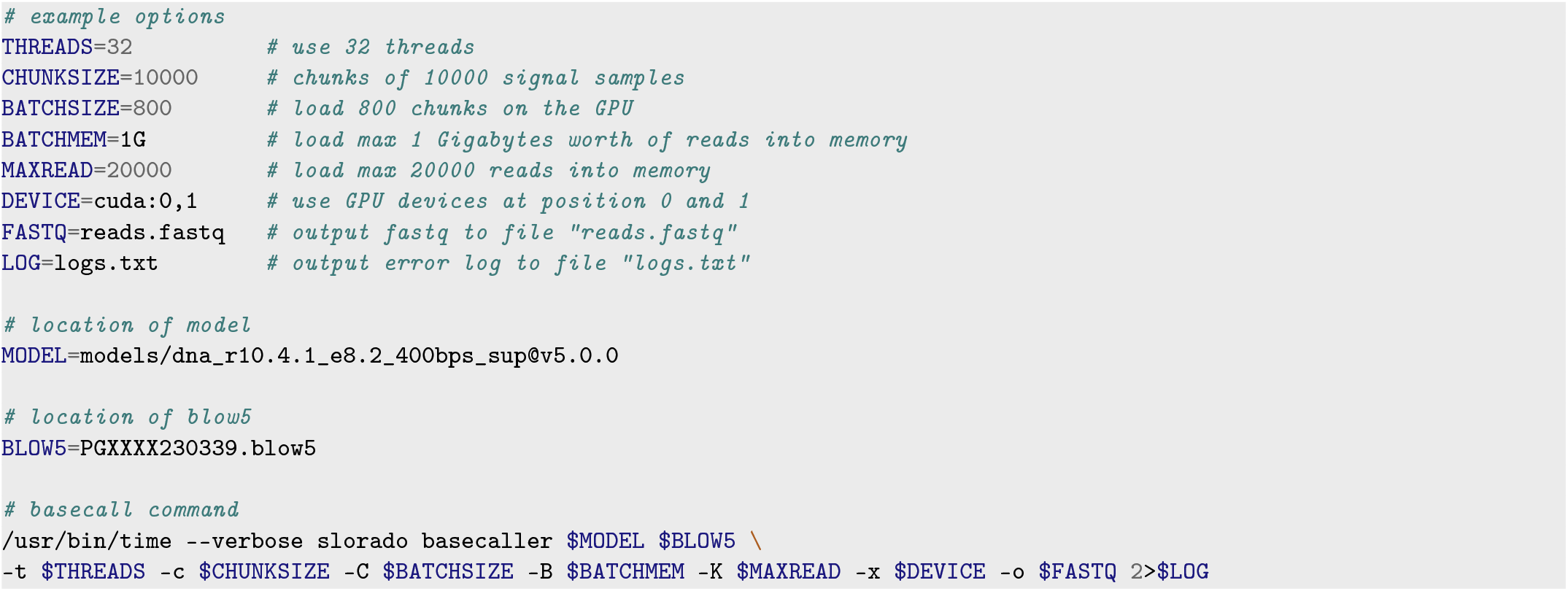

Versions:

- Slorado (except below cases): 0.4.0-beta; commit 499b1cb26875fd5c6d77029714c8c89f83cf97d3
- Slorado on NVIDIA edge device: 0.4.0-beta; commit 406ab8c104729eadbc37fcb63bfbc5cfe78058ce
- Slorado on AMD edge device: v0.4.0-beta binaries

#### Basecalling Accuracy

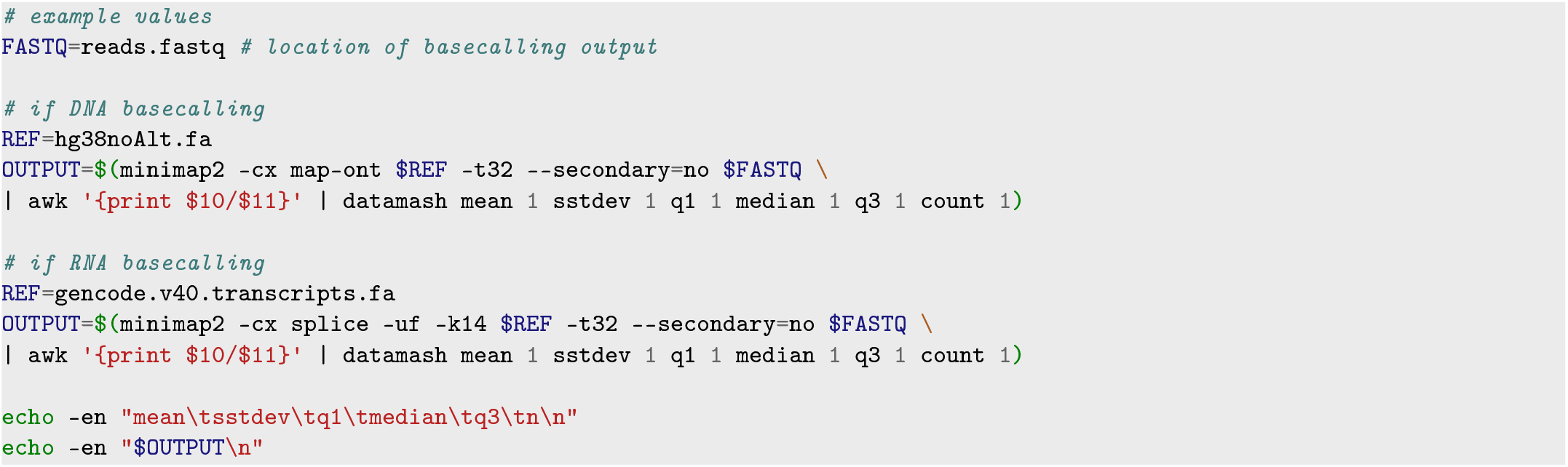

Versions:

- Datamash: v1.3.0
- Minimap2: v2.26.0

#### Multi-node scaling analysis

##### Split BLOW5

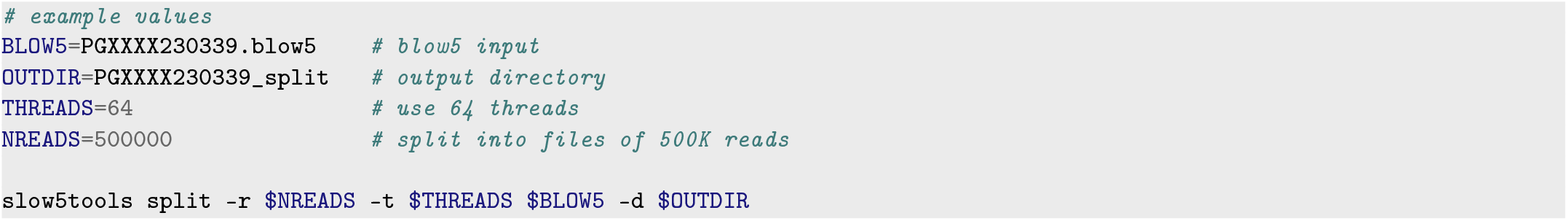

Versions:

- Slow5tools: v1.3.0

##### Launch Script

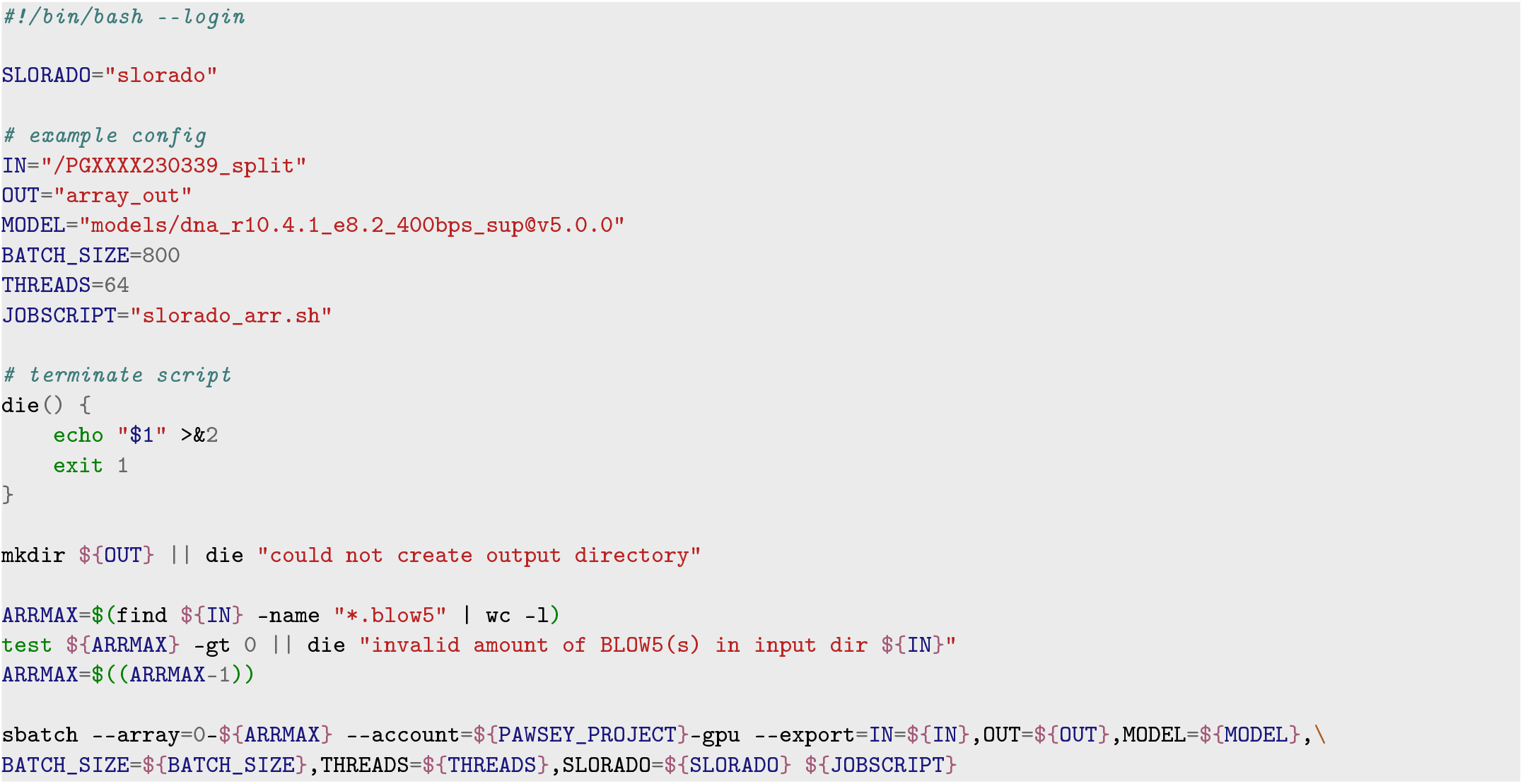

##### Job Script

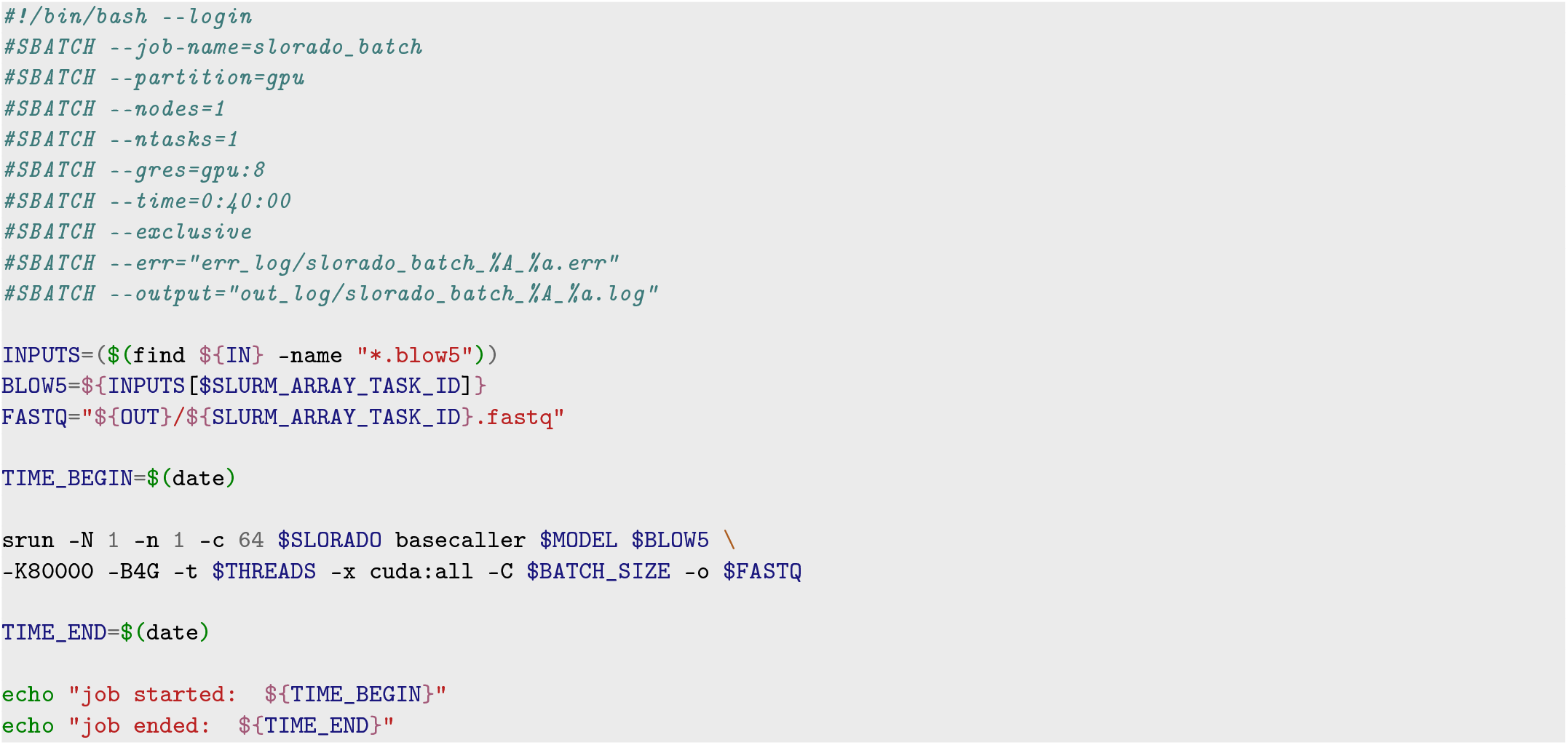

#### Dorado Basecalling

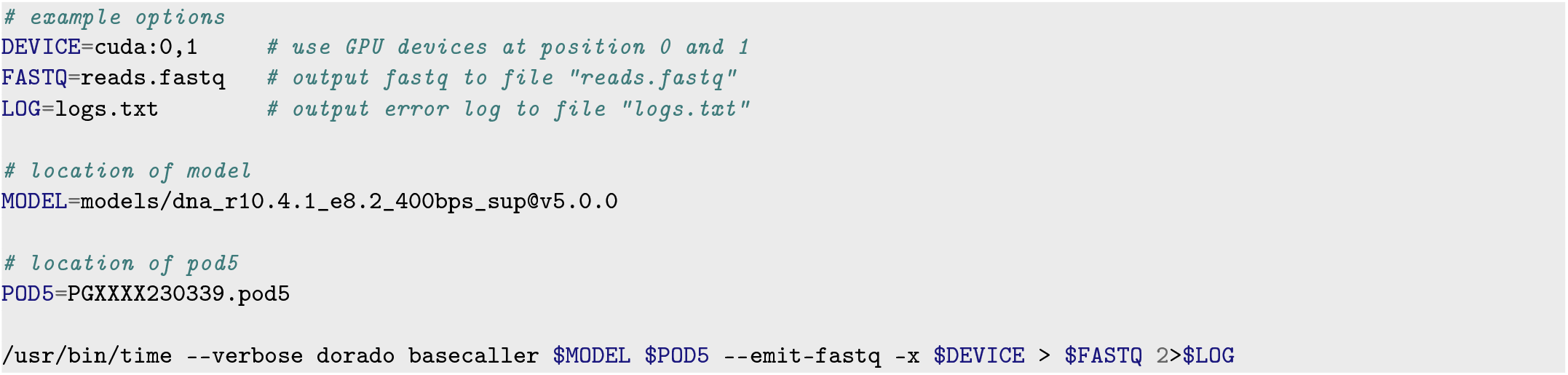

Versions:

- Dorado: v1.1.1 binaries

#### Slorado Source Code Compilation

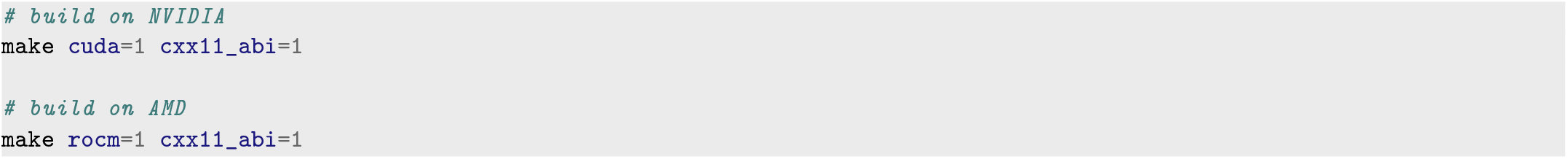

#### POD5 -> BLOW5 conversion

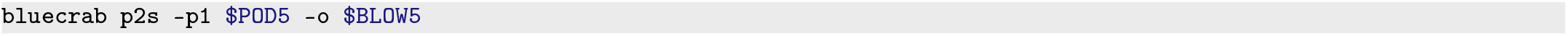

Versions:

- Bluecrab: v0.4.0

#### Variant Calling

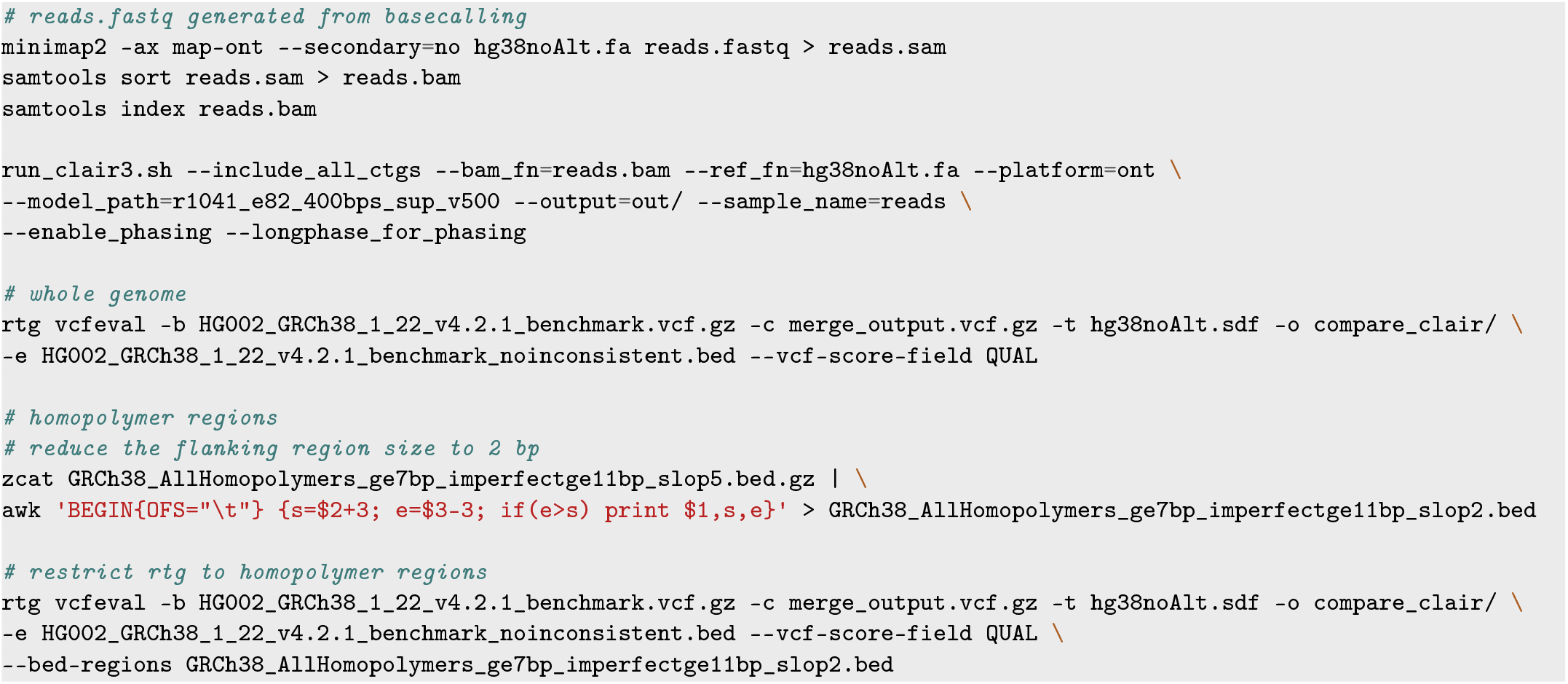

Versions:

- Minimap2: v2.26.0
- Samtools: v1.18.0
- Clair3 v1.0.10
- Rtg: v3.12.1

#### Modification Detection

##### Slorado Modification Detection

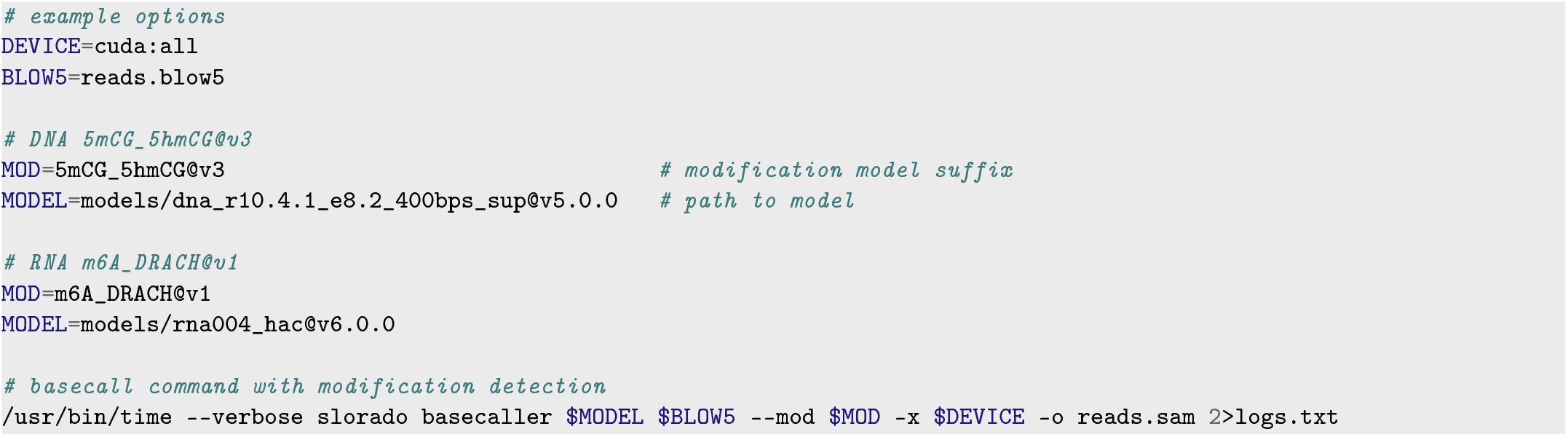

Versions:

- Slorado: v0.6.0 binaries

##### Dorado Modification Detection

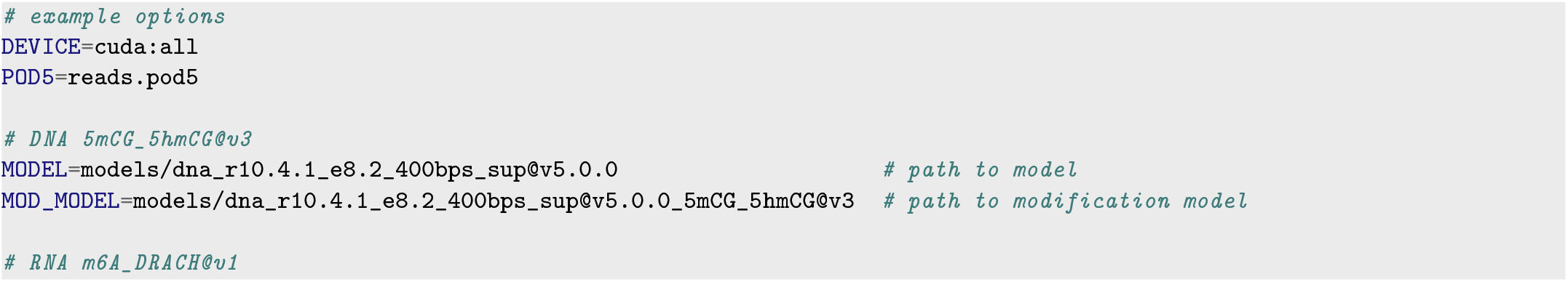
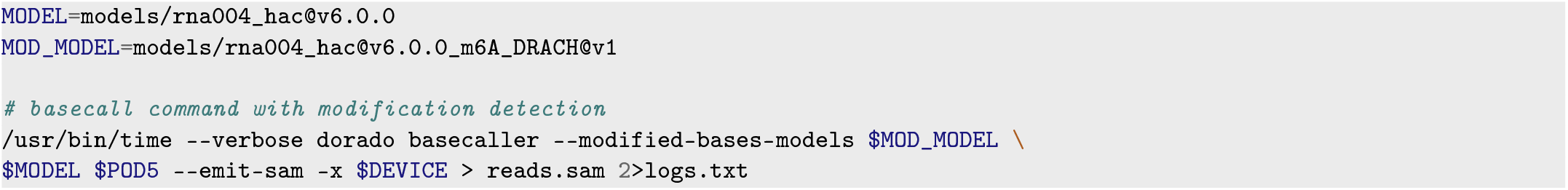

Versions:

- Dorado: v2.0.0 binaries

##### Methylation Frequency Correlation

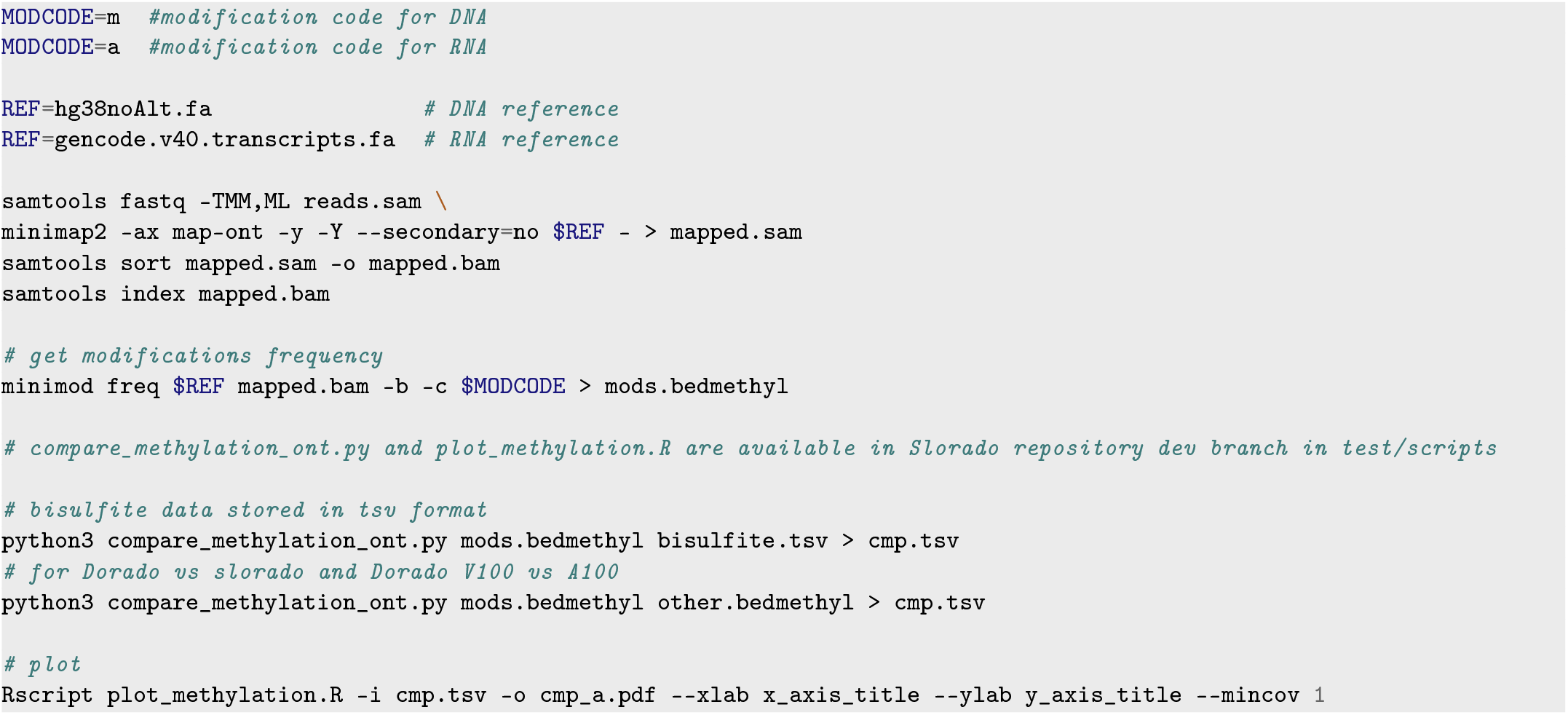

Versions:

- Minimap2: v2.26.0
- Samtools: v1.18.0
- Minimod: v0.5.0

### Supplementary Note 2 - GPU Decode subroutines

#### Algorithm S1 initialise_beam

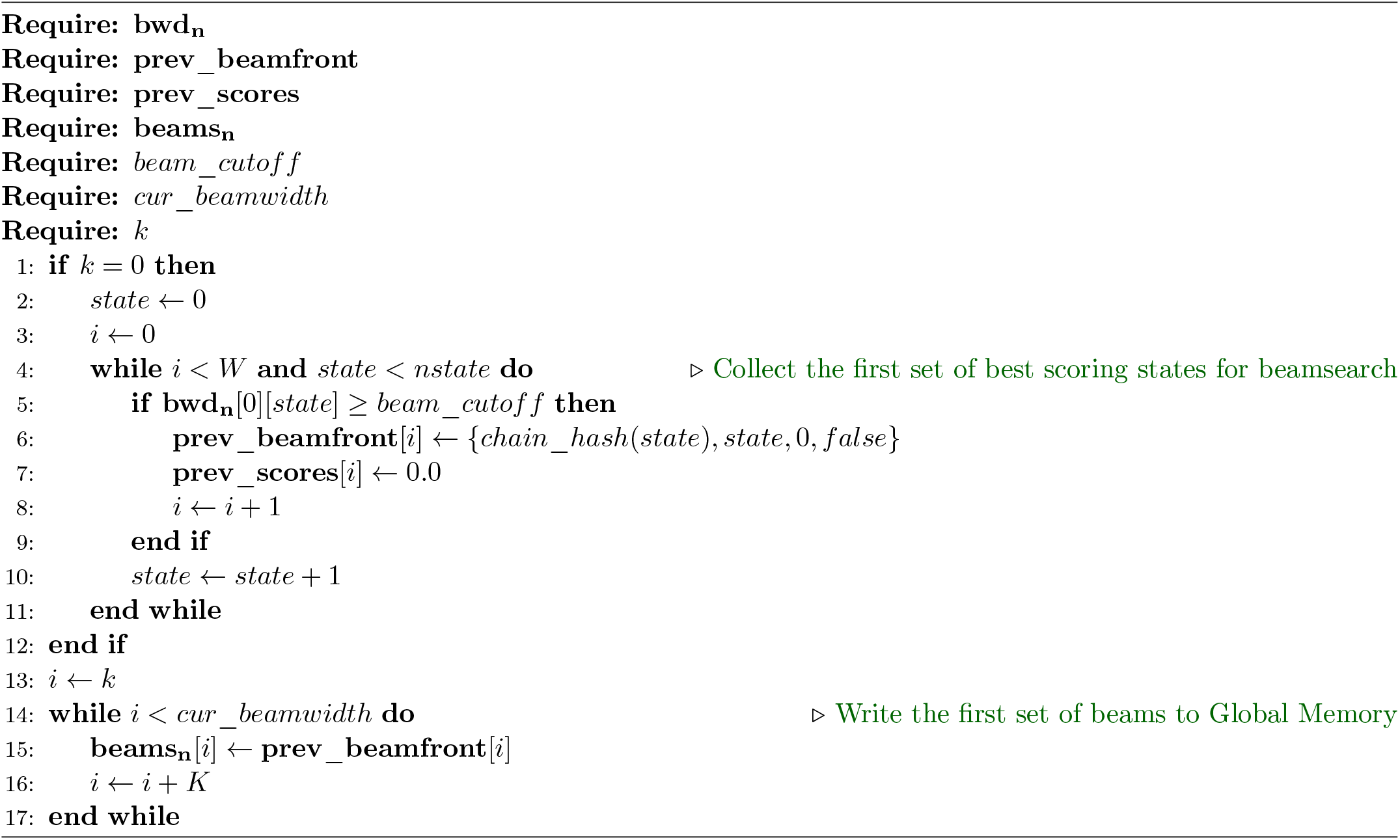

#### Algorithm S2 merge_duplicates

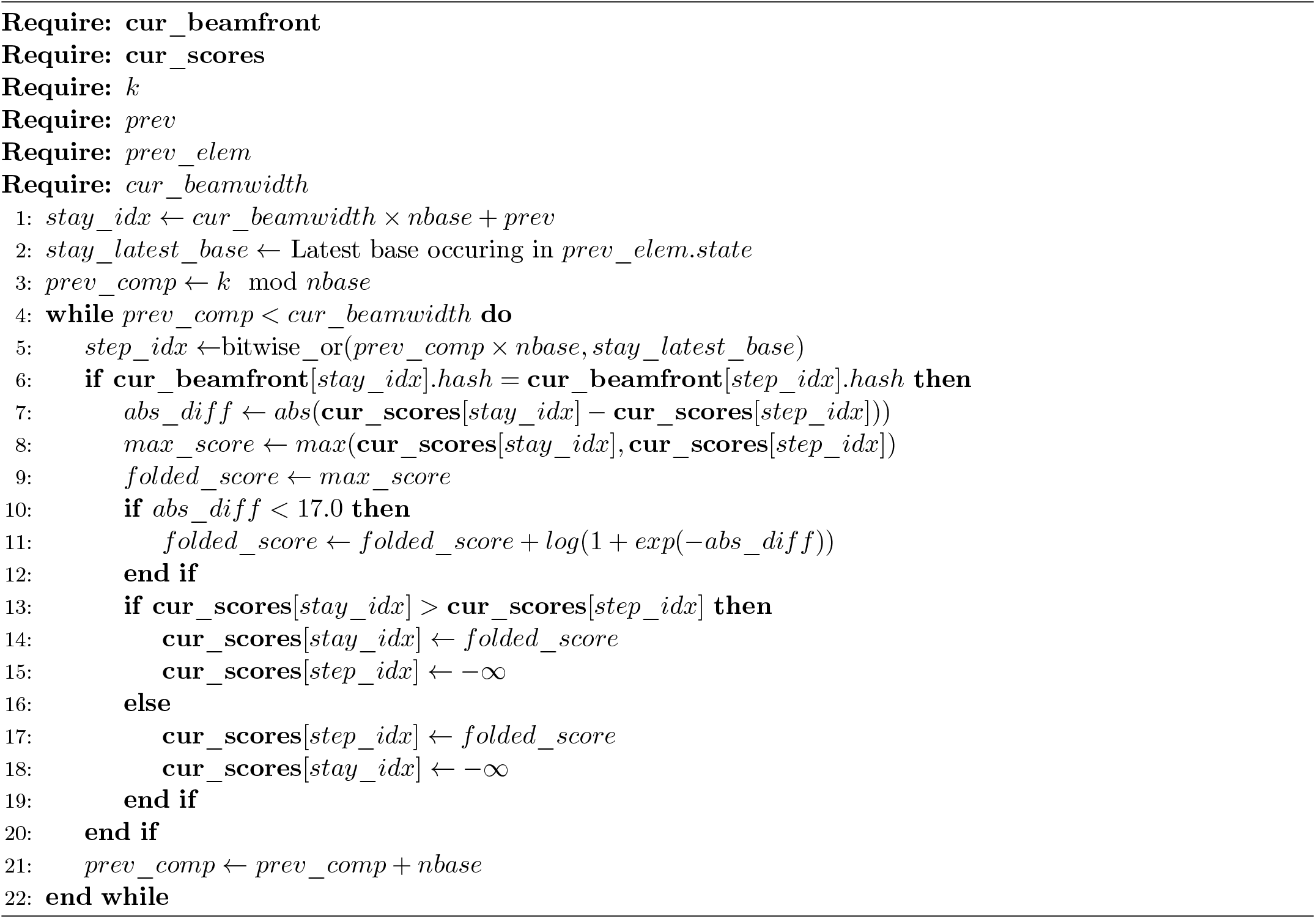

#### Algorithm S3 move_current_beam_to_previous

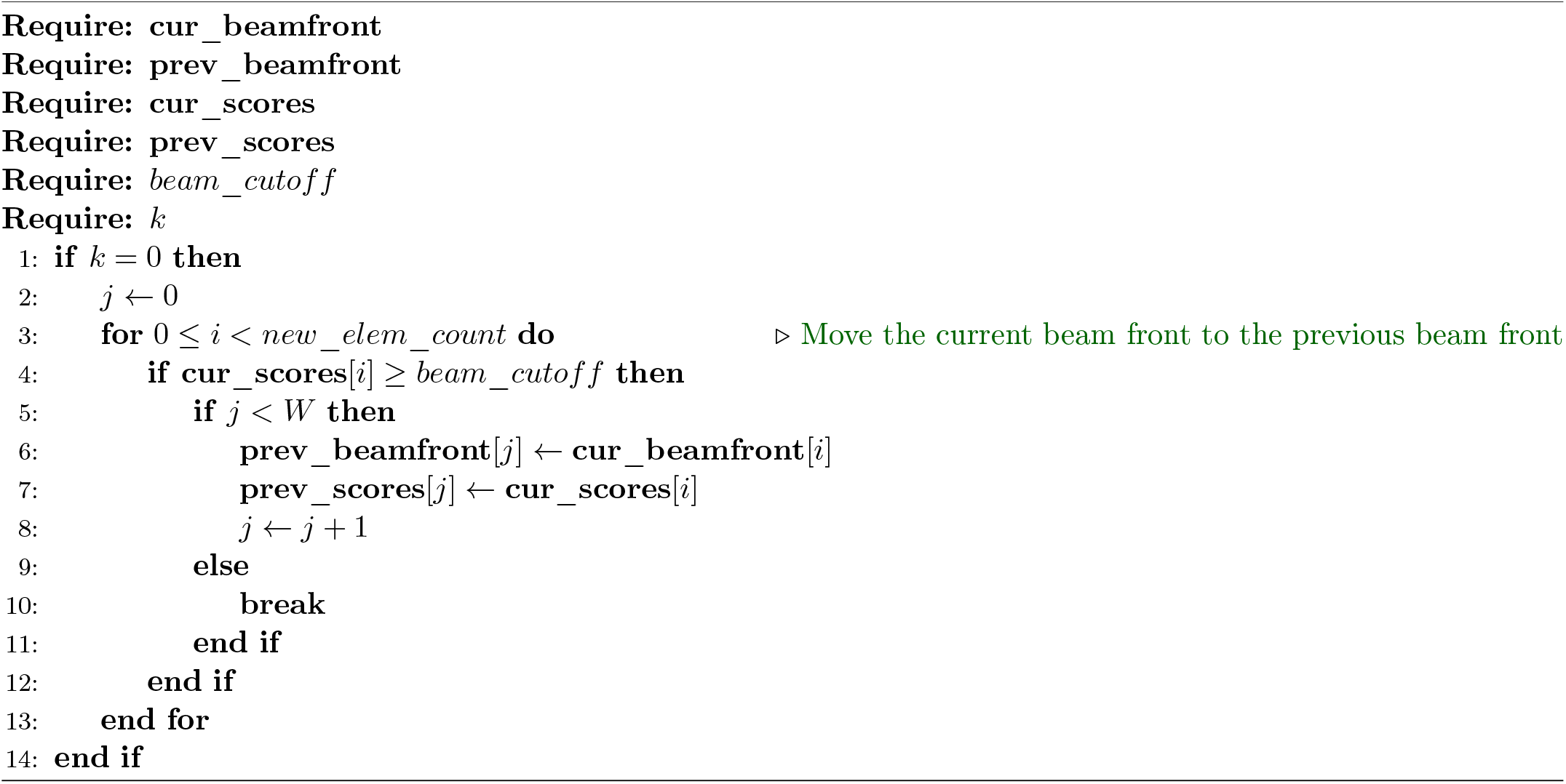

#### Algorithm S4 find_best_beam

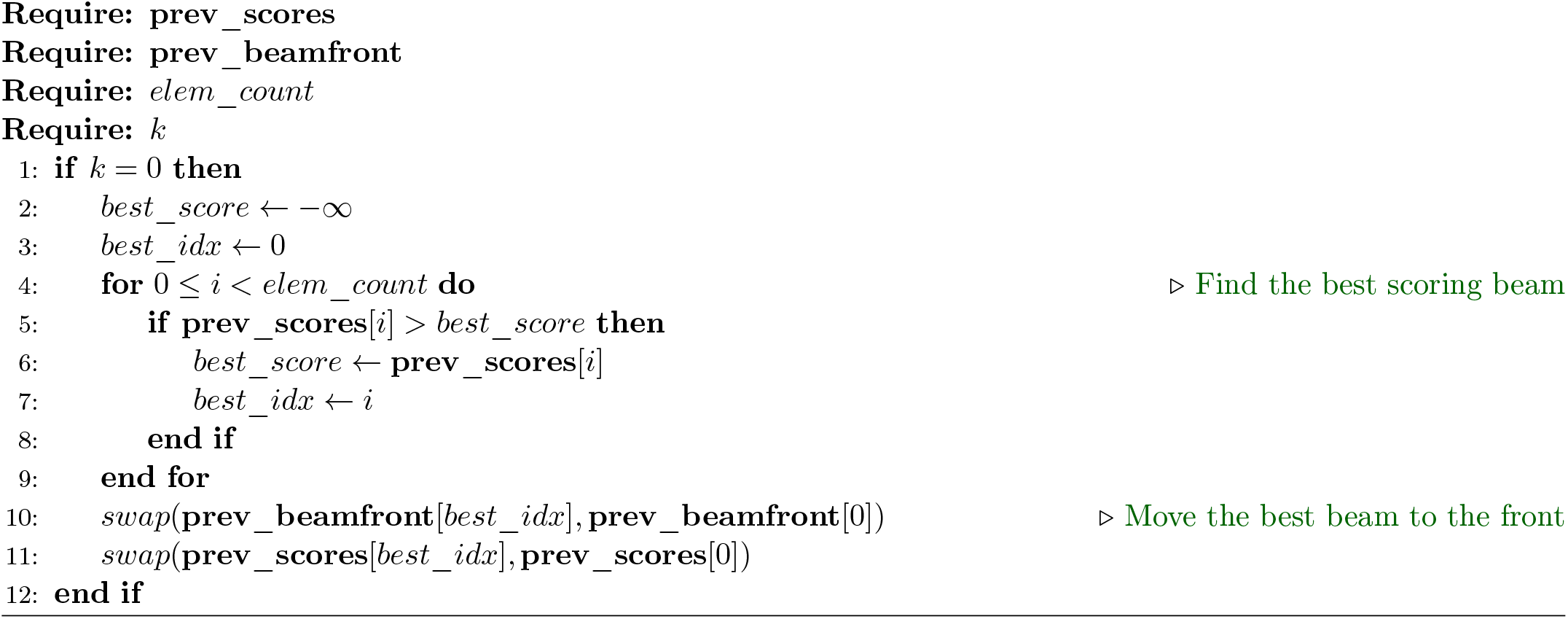

#### Algorithm S5 write_beam_results

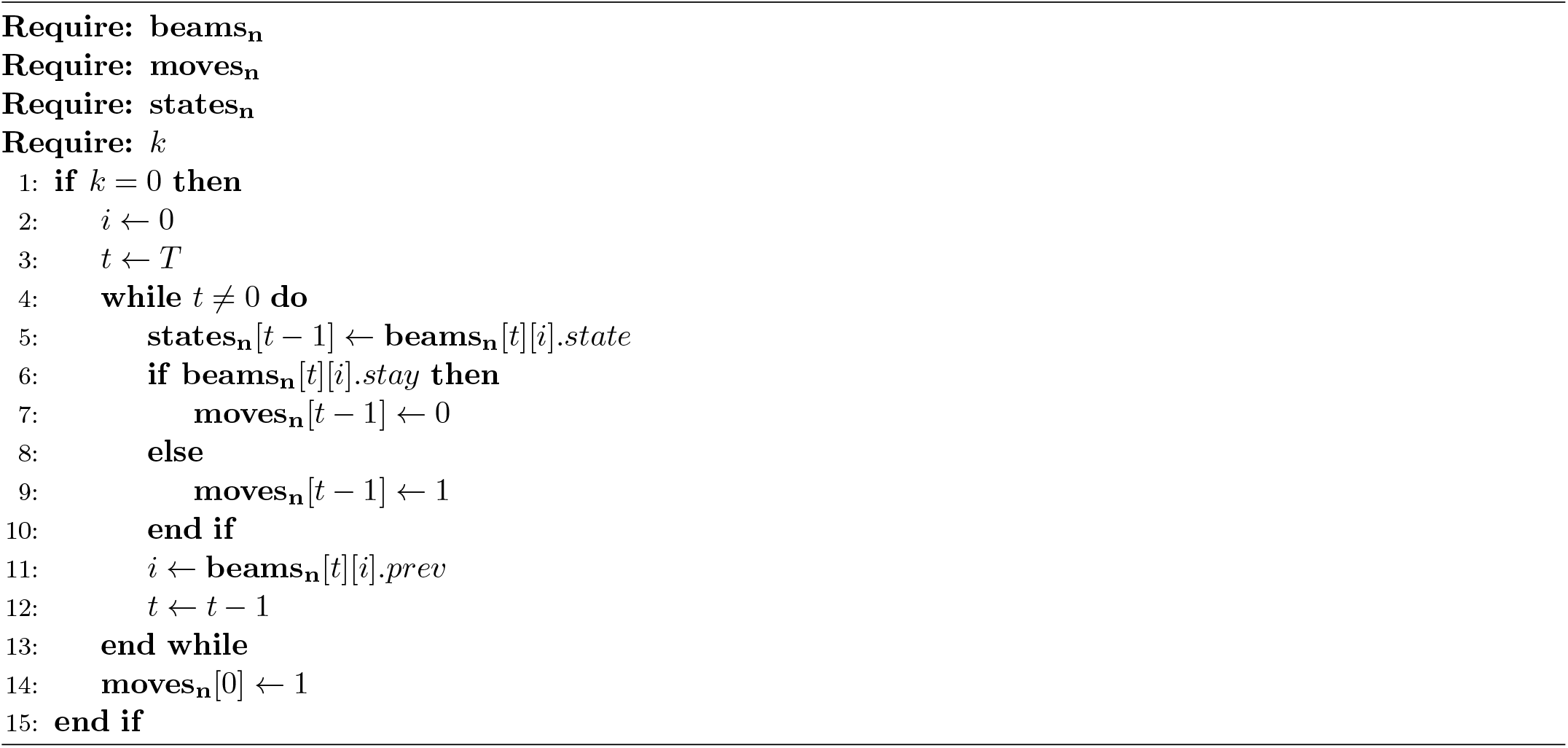

#### Algorithm S6 compute_quality

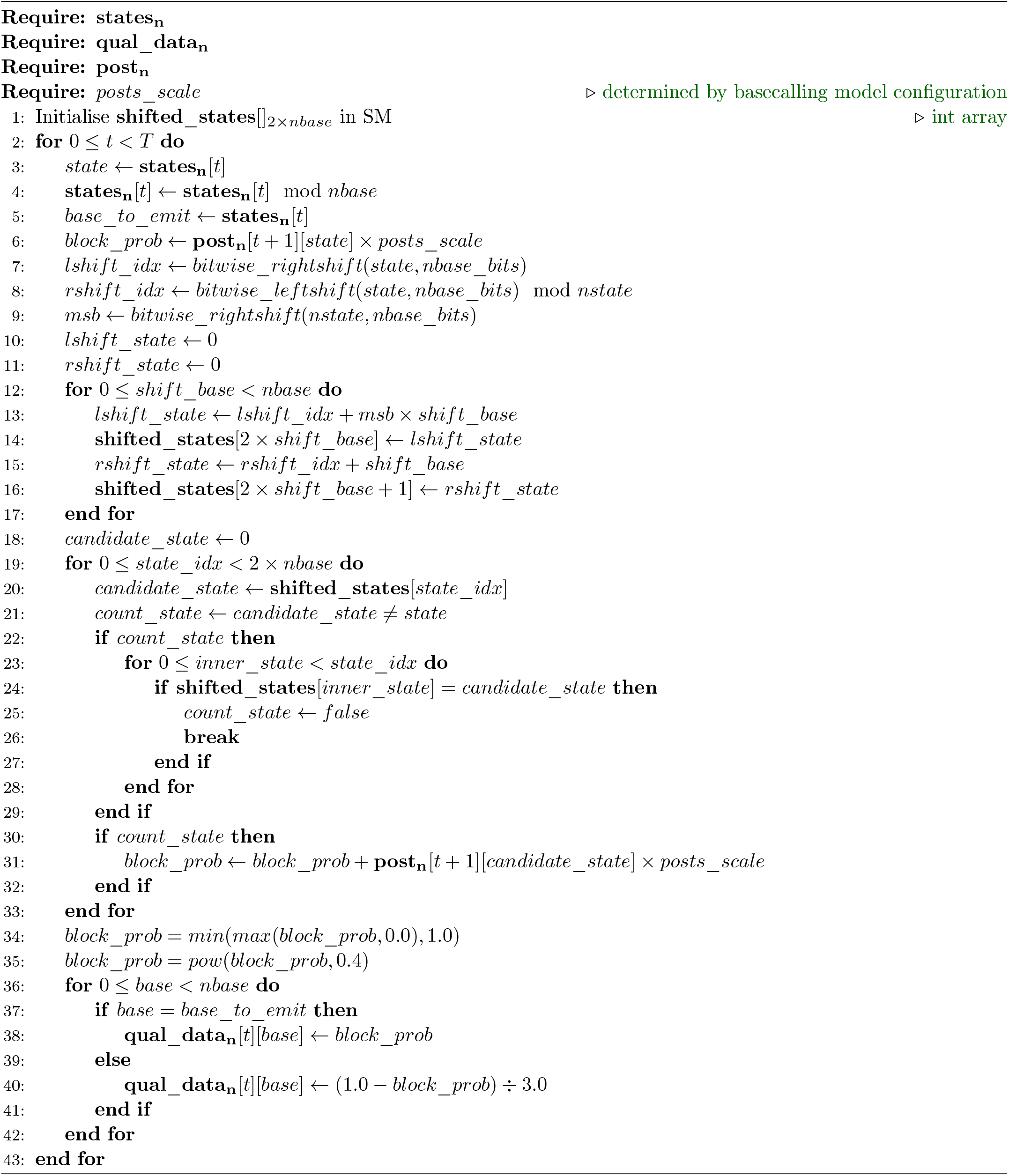

#### Algorithm S7 generate_sequence_qscore

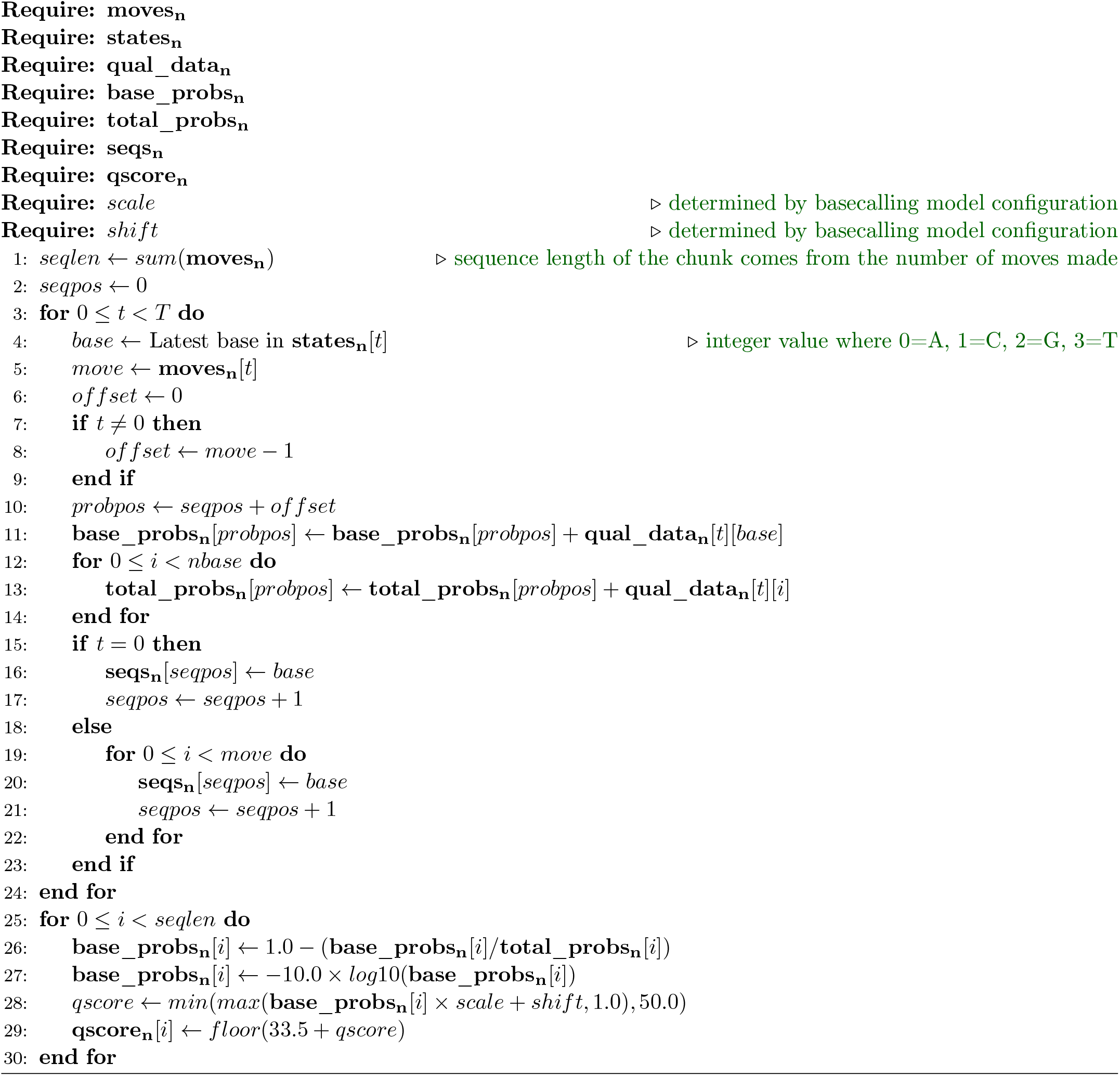

